# Multiomics approach identifies SERPINB1 as candidate progression biomarker for Spinocerebellar Ataxia type 2

**DOI:** 10.1101/2025.04.18.649103

**Authors:** Luis E. Almaguer-Mederos, J Key, NE Sen, J Canet-Pons, C Döring, D Meierhofer, Suzana Gispert-Sánchez, Dany Cuello-Almarales, Dennis Almaguer-Gotay, Lidia Matilde Osorio González, Raúl Aguilera-Rodríguez, Jacqueline Medrano Montero, Georg Auburger

**Affiliations:** Goethe University, University Hospital, Clinic of Neurology, Exp. Neurology, Frankfurt am Main, Germany; Goethe University, Medical Faculty, Dr. Senckenberg Institute of Pathology, Frankfurt am Main, Germany; Max Planck Institute for Molecular Genetics, Berlin, Germany; Center for the Investigation and Rehabilitation of Hereditary Ataxias, Holguín, Cuba

## Abstract

**Background:** Spinocerebellar ataxia type 2 (SCA2) is a polyglutamine disorder, and variants in its disease protein Ataxin-2 act as modifiers in the progression of Amyotrophic Lateral Sclerosis. There are no reliable molecular progression biomarkers for SCA2.

**Objectives:** The aim of this study was to define novel molecular progression biomarker candidates for SCA2.

**Methods:** Using cerebellar and cervicothoracic spinal cord RNA from *Atxn2*-CAG100-KnockIn and wildtype mice, a multi-omics study was conducted, followed by validation in mice and humans. Global transcriptome studies were conducted using the Clariom D microarray. Extracted proteins were analyzed by LC-MS/MS for global proteomics, and Immobilized Metal Affinity Chromatography for phosphoproteomics. Validation assessed expression by RT-qPCR, and protein abundance by quantitative immunoblots and ELISA. Patients with SCA2 were diagnosed following standard procedures, and the age at onset, SARA score, INAS count, and disease duration were used as clinical severity markers.

**Results:** Venn diagram comparisons across all OMICS datasets indicated that only *Serpinb1a*-transcript, SERPINB1A-protein and -phosphopeptides were consistently downregulated at terminal stage in 14-month-old KnockIn mice. Expression studies in cerebellum and spinal cord from 10 weeks (pre-manifest), 6-month-old (early ataxic), and 14-month-old (late ataxic stage) mice confirmed this progressive decrease at mRNA and protein level. SERPINB1 plasma levels were significantly lower in SCA2 patients, and displayed a significant association with the CAG repeat length at expanded *ATXN2* alleles and the age at onset, also showing a trend towards significance with the SARA score.

**Conclusions:** SERPINB1 was identified as novel promising biomarker with specificity for SCA2 pathomechanisms.

## Introduction

Spinocerebellar ataxia type 2 (SCA2) is a progressive neurodegenerative disorder for which there are no disease-modifying treatments. It is caused by a CAG repeat expansion mutation in the coding region of the *ATXN2* gene (1-3). CAG repeat expansions beyond 31 CAG repeats cause the typical SCA2 cerebellar syndrome, which presents with gait ataxia, dysdiadochokinesia, dysmetria, and dysarthria. This presentation is frequently accompanied by kinetic or postural tremor, leg cramps, decreased tendon reflexes, abnormal eye movements with slowed saccades, and reduced body mass index and survival (4-8). In contrast, *ATXN2* intermediate-length repeat expansions in the range of 27 to 33 units act within oligo- or polygenic backgrounds to increase the disease risk or progression of amyotrophic lateral sclerosis (ALS) (9-11), frontotemporal dementia (FTD) (12, 13), progressive supranuclear palsy (PSP) (14), and idiopathic levodopa-responsive Parkinson’s disease (15).

The *ATXN2* gene encodes Ataxin-2, a cytoplasmic RNA binding factor involved in the regulation of cytoskeletal dynamics, endocytosis, growth and cellular stress responses, mRNA translation, and the inhibition of the mechanistic target of rapamycin complex 1 (mTORC1) signaling (16-23). Upon the unstable expansion of its polyQ domain, the Ataxin-2 protein acquires a toxic gain of function leading to a multi-level atrophy, prominently of neurons but also peripheral adipocytes and central nervous system oligodendrocytes (7, 24, 25).

Though there is no curative treatment for SCA2 yet, Ataxin-2 abundance-lowering strategies, based on small chemical compounds (26-28), antisense oligonucleotides (29, 30) or Cas13 CRISPR effectors (31) against *Atxn2* mRNA, and also on pharmacological autophagy enhancement (32), proved successful in experiments conducted in murine SCA2/ALS models. Clinical trials based on those findings have been initiated. Even so, the precise assessment of experimental therapeutic interventions relies upon informative biomarkers of disease severity and progression, with a strong involvement in the pathophysiology of SCA2. Consequently, the identification of molecular events that serve as reliable biomarker surrogate endpoints is of chief significance for the development and validation of effective treatments against SCA2.

PolyQ-expanded Ataxin-2 abundance in biofluids would be an obvious candidate biomarker for SCA2, similar to what was described for huntingtin or Ataxin-3 proteins in Huntingtońs disease (HD) or SCA3, respectively (33-35). However, Ataxin-2 levels in the blood and cerebrospinal fluid are very low (https://gtexportal.org/home/gene/ENSG00000204842.14) (36), and no routine assay sensitive enough to consistently detect it in these biofluids has been developed yet (37). Nonetheless, the constitutive and ubiquitous expression pattern of Ataxin-2 increases the chance of finding easily detectable peripheral molecular biomarkers, resulting from downstream effects of the gain/loss of poly-Q expanded Ataxin-2 functions. Preliminary evidence based on studies conducted in SCA2 patients suggested the levels of the mitochondrial kinase PINK1 (38), the aggrephagy marker WDFY3 (39), trace elements (Mn, Cu, Zn, and V) (40), oxidative stress markers (41), and the serum testosterone levels (42), as candidate peripheral biomarkers for SCA2. However, most of these studies were conducted on small cohorts or lack independent validations.

More recently, the neurofilament light chain (NfL) levels in cerebrospinal fluid (CSF) or blood were proposed as a candidate biomarker in SCA2 (43-47). Likewise, elevated NfL levels were recognized in several neurologic disorders including ALS (48, 49), HD (50), and SCA1, 3 (51), and 7 (46), among others. Based on those findings, NfL levels have been used as a biomarker endpoint in clinical trials assessing the effectiveness of antisense oligonucleotides targeting *SOD1* mRNA (Tofersen) in ALS (52, 53), or targeting *HTT* mRNA (Tominersen) in HD (54, 55). Even though the levels of NfL in CSF/blood seem to be a promising biomarker in neurodegenerative disorders including SCA2, its low abundance can be detected only in few centers worldwide by costly equipment, and its low specificity together with issues regarding assays standardization complicate its use as surrogate endpoint.

Progress in high-throughput “omic” technologies provides new opportunities to understand the pathogenesis of neurodegenerative disorders (56, 57). Moreover, integrating several omics into multi-omic platforms offers further possibilities to describe relevant peripheral disease biomarkers. The use of multi-omics proved useful in biomarker research for common neurodegenerative disorders such as Alzheimeŕs (58) and Parkinsońs diseases (59), and for rare disorders, including ALS (60), and HD (61).

In the present study, a multi-omics approach was applied to an authentic SCA2 mouse model (*Atxn2*-CAG100-KIN). This revealed key pathways and molecular events linked to SCA2 pathophysiology and defined SERPINB1, a member of the SERPIN superfamily of protease inhibitors, as a candidate progression biomarker for SCA2. This finding was followed by validation studies in the *Atxn2*-CAG100-KIN mouse model and in SCA2 patients from the largest and genetically homogenous SCA2 population worldwide.

## Methods

### Study design

An initial transcriptome profiling of translation, stress granules, RNA processing, and autophago-lysosome pathway factors was followed by an unbiased multi-omics study, based on the integration of transcriptomic, proteomic and phosphoproteomic data obtained in cerebellar and spinal cord tissue from pre-manifest (10-week-old) and end-stage (14-month-old) *Atxn2*-CAG100-KnockIn, and age and gender-matched wild type (WT) mice. Validation studies were performed at transcript and protein levels in cerebellar and spinal cord tissue from pre-manifest, early- (6-month-old), and end-stage *Atxn2*-CAG100-KnockIn mice. Further validation studies were conducted in 58 patients with SCA2 and equal number of age and gender-matched healthy individuals from the same geographical region as the patients. The SCA2 patients’ cohort was enlarged to 82 individuals for correlational analyses considering SERPINB1 blood plasma levels and core markers of disease severity.

### Experimental animals

All mouse experiments were in conformity with the German Animal Welfare Act, the Council Directive of 24th November 1986 (86/609/EWG) with Annex II, and the ETS123 (European Convention for the Protection of Vertebrate Animals). All mice were housed at the Central Animal Facility (ZFE) of Goethe University Medical School, kept in individually ventilated cages with nesting material at a 12 h light/12 h dark cycle, with appropriate temperature and humidity conditions, and provided with food and water *ad libitum*. Among offspring littermates, the homozygous *Atxn2*-CAG100-KnockIn and WT animals of the same sex were selected and aged together in the same cages, for subsequent experimental group comparisons regarding gene expression, protein, and phospho-peptides abundance. WT and *Atxn2*-CAG100-KnockIn mice were genotyped as previously described (25). Both female and male mice were used during all experiments. The Regierungspräsidium Darmstadt, with approval number V54-19c20/15-FK/1083, ethically supported the study.

### SCA2 patients and healthy control individuals

The Ethics Committee of the Center for the Investigation and Rehabilitation of Hereditary Ataxias (CIRAH) approved the study protocol for SCA2 patients, and written informed consent was obtained from patients according to the Declaration of Helsinki. Blood plasma SERPINB1 levels were measured in randomly selected patients with a clinical and molecular diagnosis of SCA2, and in age and gender-matched healthy control individuals from the same geographical region as the patients. Control individuals with family history of neurodegenerative disorders were excluded from the study. Likewise, individuals with reported personal history of lung disease, Cohen syndrome, alpha 1-antitrypsin deficiency, impetigo, vasculitis, granulomatosis with polyangiitis, plasma protein metabolism disease, psoriasis, cystic fibrosis, systemic lupus erythematosus, or neutropenia, were also excluded based on their reported associations with altered SERPINB1 blood levels (https://www.genecards.org/cgi-bin/carddisp.pl?gene=SERPINB1&keywords=serpinb1#diseases; last accessed on December 20, 2024).

### Transcriptome profiling

Global transcriptomic studies based on total RNA obtained from cerebellar and cervicothoracic spinal cord tissue from three WT and equal number of end-stage (14-month-old) homozygous *Atxn2*-CAG100-KnockIn mice were conducted as previously described (Canet-Pons et al., 2021). Briefly, the total RNA integrity was determined using the 2100 Bioanalyzer with the Nano Assay (Agilent Technologies, Santa Clara, CA). Single−stranded cDNA (ss−cDNA) was generated using the GeneChipTM WT PLUS Reagent Kit (Applied Biosystems, Foster City, CA) following the manufacturer’s instructions. The ss−cDNA was fragmented and labeled immediately before the hybridization to a Clariom D Array (Thermo Fisher Scientific). The arrays were scanned with the Affymetrix GeneChip Scanner, and the resulting data were processed with the Transcriptome Analysis Console (TAC) 4.0.1 (Applied Biosystems) using default software parameters.

For data processing, a list of 720 factors was initially obtained by filtering the transcriptome dataset in the Transcriptome Analysis Console for protein translation-related factors. In addition, a list of 505 factors that defines an up-to-date manually curated catalog of factors involved in translation was obtained from the Proteostasis Consortium (https://www.proteostasisconsortium.com/pn-annotation; last accessed on February 4, 2025). Next, for the analysis of stress granules and RNA processing factors, a comprehensive list of 967 RNA processing (GO: 0006396) factors was obtained from the Mouse Genome Informatics (MGI) resource (https://www.informatics.jax.org; last accessed on February 4, 2025) (74). In addition, a list of 407 factors that defines an up-to-date manually curated catalog of stress granules proteins was obtained from the RNAgranule database (version 2.0) (http://rnagranuledb.lunenfeld.ca; last accessed on February 4, 2025) (75). Afterwards, for the analysis of dysregulated factors with prominent roles in the Autophagy-Lysosome pathway (ALP), a list of 870 factors that defines an up-to-date manually curated catalog of translation factors was obtained from the Proteostasis Consortium (https://www.proteostasisconsortium.com/pn-annotation; last accessed on February 4, 2025).

### Proteome profiling

Cerebellar and spinal cord protein extracts from eight WT and six end-stage (14-month-old) homozygous *Atxn2*-CAG100-KnockIn mice, and cerebellar protein extracts from three WT and five pre-manifest (10-week-old) homozygous *Atxn2*-CAG100-KnockIn mice were used for proteome profiling. In short, tissue samples were homogenized under denaturing conditions, and then boiled at 95 °C for 10 min, sonicated for 10 min, and centrifuged at 16,000 rcf for 10 min at 4 °C. Supernatants were transferred into new protein low binding tubes (Eppendorf, Germany). Lysed and trypsin-digested samples were desalted over C18 columns, reconstituted in 2% formic acid in water, and separated by strong cation exchange chromatography (SCX, 3M Purification, Meriden, CT). Eluates were first dried in a SpeedVac, then dissolved in 5% acetonitrile and 2% formic acid in water, briefly vortexed, and sonicated in a water bath for 30 sec previous to injection to nano-LC-MS/MS. The LC-MS/MS was carried out by nanoflow reverse-phase liquid chromatography (Dionex Ultimate 3000, Thermo Scientific) coupled online to a Q-Exactive HF Orbitrap mass spectrometer (Thermo Scientific), as previously reported (62). Raw MS data were processed with MaxQuant software (v2.1.0.0) and searched against the mouse proteome database UniProtKB with 55,366 entries, released in March 2021. Proteomics data were filtered to include only proteins detected in at least three replicate per genotype. Fold-changes and p-values were computed independently for each tissue.

### IMAC Phosphoproteomics

Mouse cervicothoracic spinal cord tissue was obtained after cervical dislocation from four WT and equal number of end-stage (14-month-old) homozygous *Atxn2*-CAG100-KnockIn mice, then immediately frozen and transported in liquid nitrogen. Phosphoproteome profiling of spinal cord tissue was performed at Cell Signaling Technology INC. using their PTMScan and Fe-IMAC services, as previously described (16). Searches were conducted against the most recent update of the Uniprot *Mus musculus* database with a mass accuracy of +/-20 ppm for precursor ions and 0.02 Da for productions. Results were filtered with mass accuracy of +/– 5 ppm on precursor ions and the existence of the intended motif.

### Quantitative reverse transcription polymerase chain reaction in spinal cord, cerebellum, and fibroblasts

Total RNA isolation from cerebellar and spinal cord tissue was performed with TRIzol Reagent (Sigma Aldrich, USA) according to manufacturer’s instructions. Total RNA yield and purity were quantified using a Tecan Spark plate reader (Tecan Group Ltd, Switzerland) at 230, 260, and 280 nm, in a NanoQuant plate. cDNA synthesis was performed from 1 μg of total RNA template using the SuperScript IV VILO kit (Invitrogen, USA) according to the manufacturer’s instructions. Gene expression profiles were assessed by quantitative reverse transcription polymerase chain reaction (RT-qPCR) using a StepOnePlus^TM^ (96 well) Real-Time PCR System (Applied Biosystems, USA). RT-qPCRs were run in technical duplicates on cDNA from 25 ng total RNA, with 1 μl TaqMan® Assay, 10 μl FastStart Universal Probe Master 2× (Rox) Mix (Roche, Switzerland) and ddH_2_O up to 20 μl of total volume. The PCR cycling conditions were 50 °C for 2 min, 95 °C for 10 min, followed by 40 cycles of 95 °C for 15 min and 60 °C for 1 min. The gene expression TaqMan® assays (Thermo Fisher Scientific, Waltham, Massachusetts, USA) used for this study were: *Ctsb* (Mm01310506_m1), *Ctsd* (Mm00515586_m1), *Ctsl* (Mm00515597_m1), *Ctss* (Mm01255859_m1), *Ctsz* (Mm00517697_m1), *Klk6* (Mm00478322_m1), *Serpina3n* (Mm00776439_m1), *Serpinb1a* (Mm01610780_m1), *Serpind1* (Mm00433939_m1), and *Serpini1* (Mm00436740_m1). The data were analyzed via the 2^−ΔΔCt^ method (63), using *Tbp* (Mm00446973_m1) as housekeeping gene.

### Protein extraction and immunoblots

Tissue samples from mouse cerebellum and cervicothoracic spinal cord sections were homogenized with a motor pestle in 5-10× weight/volume amount of RIPA buffer consisting of 50 mM Tris-HCl (pH 8.0), 150 mM NaCl, 2 mM EDTA, 1% Igepal CA-630 (Sigma Aldrich, USA), 0.5% sodium deoxycholate, 0.1% SDS, cOmplete™ Protease Inhibitor Cocktail (Roche, Switzerland), and Halt™ Phosphatase Inhibitor Cocktail (Thermo Fisher Scientific, Inc., USA). The resulting protein suspensions were sonicated, and protein concentration was determined in a Tecan Spark plate reader (Tecan Group Ltd, Switzerland) using a Pierce™ BCA protein assay kit (Thermo Fisher Scientific, Inc., USA). 15 to 25 μg of total proteins were mixed with 2× loading buffer consisting of 250 mM Tris-HCl pH7.4, 20% glycerol, 4% SDS, 10% 2-mercaptoethanol, and 0.005% bromophenol blue, incubated at 90 °C for 5 min, separated on 8-15% polyacrylamide gels at 120 Volts, and transferred to nitrocellulose membranes (0.2 µm) (Bio-Rad Laboratories, Inc., USA). The nitrocellulose membranes were blocked in 5% BSA/TBS-T, and incubated overnight at 4 °C with primary antibodies. Afterwards, the nitrocellulose membranes were incubated for 1 h at room temperature, with fluorescently labeled secondary IRDye® 800CW goat anti-mouse (LI-COR 926-32210, 1:10,000), IRDye® 800CW goat anti-rabbit (LI-COR 926-32211, 1:10,000), IRDye® 680RD goat anti-mouse (LI-COR 926-68070, 1:10,000) or IRDye® 680RD goat anti-rabbit (LI-COR 926-68071, 1:10,000). Membranes were scanned using an Odyssey® Classic Imager. Image visualization and quantification of signal intensities was performed using Image Studio^TM^ software (version 5.2) (LI-COR Biosciences, Ltd., UK). The following primary antibodies were used: KLK6 (Invitrogen PA5−86829, 1:1000), and SERPINB1A (Invitrogen PA5−76875, 1:1000). ACTB (Sigma A5441, 1:10000) or GAPDH (Calbiochem CB1001, 1:10000) served as loading controls.

### Quantification of SERPINB1 plasma levels

The quantification of SERPINB1 plasma levels was done in duplicates using the Human SERPINB1 (Sandwich ELISA) ELISA Kit - LS-F13273 (LS Bio; Seattle, Washington, USA), following manufacturer’s instructions. A four-parameter logistic fitted standard curve for calculating the SERPINB1 plasma levels was generated from the Arigo Biolaboratories website (https://www.arigobio.cn/ELISA-calculator; last accessed on February 17, 2025).

### Clinical and genetic assessments in SCA2 patients

The clinical diagnosis of SCA2 was based on the presence of gait ataxia, dysarthria, dysmetria, dysdiadochokinesis, dysphagia and slow saccades. The age at onset was defined as the onset of motor impairment, and it was ascertained through interviews with patients and close relatives, or the review of patients’ medical records. The Scale for the Assessment and Rating of Ataxia (SARA) score was used as a measure of ataxia severity. This scale ranges from zero to 40 points, increasing with cerebellar disease progression (64). In addition, the Inventory Non-Ataxia Symptoms (INAS) was applied as a measure of disease severity in terms of extracerebellar involvement (65). Additional information regarding age, gender, and disease duration in years from disease onset to latest examination, was retrieved. Disease stage was assessed as previously specified (66).

### Bioinformatics analyses

STRING web-server (https://string-db.org/) (version 11.0, last accessed on March 4, 2025) (67) was used for pathway enrichment analyses using (i) lists of dysregulated transcripts identified in the *Atxn2*-CAG100-KnockIn cerebellar and spinal cord transcriptome profiling, or (ii) lists of commonly dysregulated factors in the *Atxn2*-CAG100-KnockIn cerebellar and spinal cord transcriptomes and proteomes, and in the spinal cord phosphoproteome, as input. Different functional classification frameworks were used, including Gene Ontology (GO), Kyoto Encyclopedia of Genes and Genomes (KEGG) Pathways, Reactome Pathways, and The Mammalian Phenotype Ontology (Monarch). Venn diagrams created using the Venn diagram web tool from the University of Ghent, Ghent, Belgium (http://bioinformatics.psb.ugent.be/webtools/Venn; last accessed on February 28, 2025), were used to identify factors commonly dysregulated across omic datasets. The PhosphoSitePlus® database was searched for dysregulated phospho-sites identified in the *Atxn2*-CAG100-KnockIn mice spinal cord phospho-proteome profiling (68) (www.phosphositeplus.org; last accessed on March 10, 2025).

### Statistical analyses

Unpaired Student t-tests with Welch’s correction were used to establish comparisons for continuous variables between homozygous *Atxn2*-CAG100-KIN and WT animals, and between SCA2 patients and control individuals. Chi-square test was applied to look for differences in gender distributions between SCA2 patients and control individuals. Normal distribution was verified for all variables using the Kolmogorov–Smirnov test. Correlations between blood plasma SERPINB1 levels and markers of disease severity in the SCA2 cohort were established by using the Pearson correlation test. The age at onset was corrected for the CAG repeat length at expanded *ATXN2* alleles by simple linear regression. Bar charts depicting the mean and standard error of the mean (SEM) values, and a correlation matrix diagram were used for data visualization. All statistical analyses were conducted using GraphPad Prism software (version 8.4.2) (GraphPad Software Inc., USA). Significance was assumed at p<0.05 and highlighted with asterisks: Trend (T)<0.01; *p<0.05, **p<0.01, ***p<0.001, ****p<0.0001.

## Results

### Transcripts encoding translation factors are predominantly upregulated since early age

In light of the relevance of Ataxin-2 as a translation regulation factor (18, 19, 62, 63), we first explored the cerebellar and spinal cord transcriptomes of pre-manifest and end-stage KnockIn mice looking for significant dysregulations in translation factors.

Only *Eif5a2* transcript downregulation was consistent among the four panels of Figure S1, but reflects progression only in spinal cord and poorly (Tables S1, S2). The upregulation of ribosomal factors is reminiscent of the *Atxn2*-KO effect (62).

Major progressive increases in transcript expression levels involved several pseudogenes for ribosomal proteins, with fold change differences (FC diff.) ranging from 0.53 for *Rps25-ps1* to 0.79 for *Rpl10a-ps1*. Furthermore, transcripts encoding the Poly (A) Binding Protein Cytoplasmic 1 (*Pabpc1*, FC diff. 0.65), and the Eukaryotic Translation Initiation factors 2 Alpha Kinase 2 (*Eif2ak2*, FC diff. 0.64), and 2 subunit 3, Y-linked *(Eif2s3y*, FC diff. 0.69), showed progressive upregulations. Conversely, mRNAs encoding the Eukaryotic Translation Initiation factors 5A2 (*Eif5a2*, FC diff. -0.59), and the UTP14B Small Subunit Processome Component (*Utp14b*, FC diff. -0.88), showed major progressive decreases in their expression levels (Table S2).

### Stress granules and RNA processing transcripts dysregulations are prominent since early age

Given the relevance of Ataxin-2 as an RNA processing factor and as a component of stress granules (23, 64-66), we next surveyed additional stress granule and RNA processing factors as a strategy to define the relative contributions of these cellular processes to SCA2 pathophysiology.

Altogether, progressive induction of RNA processing factors was prominent in end-stage spinal cord, but no consistently altered biomarker was identified (Figure S2; Table S3).

### Transcripts encoding factors involved in the Autophagy-Lysosome pathway are induced

Because of the involvement of polyQ-expanded Ataxin-2 with the receptor-endocytosis apparatus and macroautophagy (16, 21, 32, 67), we examined the transcriptomes looking for dysregulated factors with prominent roles in the Autophagy-Lysosome pathway (ALP).

Among all ALP factors examined, transcripts encoding lysosomal cathepsins S, Z, and L were prominently upregulated (Figure S3, Table S4). Validation efforts confirmed major upregulations of cathepsins encoding transcripts in the cerebellum and spinal cord of end-stage KnockIn mice (Figure S4). Nonetheless, only the *Kif5b* transcript showed significantly changed expression levels in cerebellar and spinal cord tissues.

Overall, the autophagy-lysosome pathway is progressively induced in both affected tissues, but *Kif5b* as single consistently altered component did not show very strong decreases.

### Kallikrein 6 (Klk6) is prominently downregulated since early age

Interestingly, inspection of factors involved in cellular proteolytic breakdown in addition to cathepsins allowed identification of *Klk6* transcripts as progressively downregulated in cerebellar and spinal cord tissue of KnockIn mice. Cerebellar transcriptomes revealed a -2.11 and -3.33−fold reduced *Klk6* expression levels in the pre-manifest and end-stage mice, respectively. Even more pronounced *Klk6* downregulations were found in the spinal cord, with fold changes of -4.03 and -23.08 in the pre-manifest and end-stage mice, respectively.

Validation studies confirmed the progressive significant downregulation of *Klk6* transcript, with reductions to 58.4% in the pre-manifest, 26.6% in the early ataxic stage, and 12.1% in the end-stage KnockIn mice cerebellum. Similarly, reductions of *Klk6* transcript expression levels to 38.3% in the pre-manifest, 22.3% in the early ataxic stage, and 21.2% in the end-stage KnockIn mice spinal cord were observed. Quantitative immunoblots produced similar results at the protein level in end-stage KnockIn mice, with reduced KLK6 abundance to 27.5% in the cerebellum and 32.1% in the spinal cord (Figure S5). Altogether, these results support KLK6/*Klk6* as a candidate progression biomarker for SCA2.

### Multi-omics data analysis define commonly dysregulated factors

After transcriptomes profiling we proceeded to integrate analyses of our transcriptomic, proteomic, and phosphoproteomic datasets. We initially conducted Venn diagrams analyses to define commonly dysregulated factors in the cerebellar and spinal cord transcriptomes and cerebellar proteome of pre-manifest KnockIn mice. As a result, no upregulated factor with nominal significance and a ≥1.5−fold change was shared across all three datasets. There were three factors commonly downregulated across all omics datasets: *Serpinb1a*, *Mobp* encoding the Myelin Associated Oligodendrocyte Basic Protein, and *Sema7a* which codes for a member of the Semaphorin family of proteins involved in dendrite growth regulation and recently implicated in ataxia (68) (Figure S6-A, B).

No upregulated factor was shared across all five datasets of end-stage KnockIn mice (Figure S6-C). STRING analysis of the 17 factors commonly upregulated in at least three out of five datasets produced significant enrichments for “Positive regulation of neuron projection development” (GO: 0010976, FDR 0.0119) and “Lysosome” (GOCC: 0005764, FDR 0.0083). On the other hand, only SERPINB1A/*Serpinb1a* was commonly downregulated across all spinal cord omics datasets (Figure S6, D). STRING analysis of the 26 factors commonly downregulated in at least three of five datasets produced significant enrichments for “Synapse” (GO: 0045202, FDR 0.009), among others.

Jointly, consistency analyses across transcriptome, proteome and phosphoproteome profiles with progression from pre-manifest to end-stage identified SERPINB1A as single most promising biomarker, with massive downregulations.

### Omics datasets provide evidence for significant dysregulations among several members of the serpin superfamily of protease inhibitors

SERPINB1A belongs to the serpin superfamily of protease inhibitors that target cathepsins and kallikrein-related peptidases. Given that cathepsins showed prominent upregulations in the transcriptomes of end-stage KnockIn mice and that there is a progressive downregulation of *Klk6* since early age, we wondered if additional serpin family members showed dysregulations that may explain protease expression level/abundance changes resulting from Ataxin-2 polyQ-expansion.

Detailed examination of the KnockIn mice transcriptomes revealed several dysregulated serpins reaching nominal significance in addition to *Serpinb1a*. Most of these serpins were downregulated in every tissue and age group examined (Figure S7; Table S5). Validation studies confirmed major serpin transcriptional dysregulations (Figure S8).

Overall, the multi-omics data indicate that the dysregulations of serpins may be revealing compensatory cellular efforts to promote the efficient degradation of polyQ-expanded Ataxin-2 aggregates by suppressing inhibitory actions of serpins on the activity of relevant proteases, with a prominent role for glial cells. SERPINB1A downregulation is clearly prominent.

### *SERPINB1A* protein and Serpinb1a transcript are consistently downregulated

*Serpinb1a* transcript showed -2.20 and -3.47−fold reduced expression levels in the cerebellar and spinal cord transcriptomes of pre-manifest 10-week-old KnockIn mice, and expression level fold changes of -2.36 and -3.03 in the cerebellar and spinal cord transcriptomes of end-stage KnockIn mice. Notably, results from the cerebellar proteome of pre-manifest KnockIn mice showed a significantly 1.96−fold (p=0.0002) reduced abundance for SERPINB1A, and similar results were apparent in the cerebellar (FC 3.73, p=0.0003) and spinal cord (FC 2.06, p=0.0001) proteomes of end-stage mice. Besides, two significant hypophosphorylations emerged in the spinal cord phosphoproteome (4.1−fold at Ser62, 5.1−fold at Ser300), probably corresponding to the reduced protein abundance of SERPINB1A.

Validation efforts confirmed the progressive reduction in *Serpinb1a* transcript and SERPINB1A protein abundance in 10-week-old (pre-manifest), six-month-old (early ataxic stage), and 14-month-old (late ataxic stage) KnockIn mice (Figure 1). In the cerebellum, SERPINB1A protein abundance reductions to 58.2, 57.2, and 29.7% were apparent in the pre-manifest, early, and late ataxic disease stages, respectively. Corresponding figures at the transcript level were 66.9, 34.5, and 25.8%, respectively. Similarly, reductions of SERPINB1A protein abundance in the spinal cord of pre-manifest, early, and late ataxic stage mice were 50.8, 42.3, and 40.9%, respectively. Likewise, *Serpinb1a* transcript was reduced in the spinal cord to 38.2% in the pre-manifest, 30.6% in the early ataxic stage, and 15.4% in the late ataxic stage KnockIn mice.

**Figure 1.**
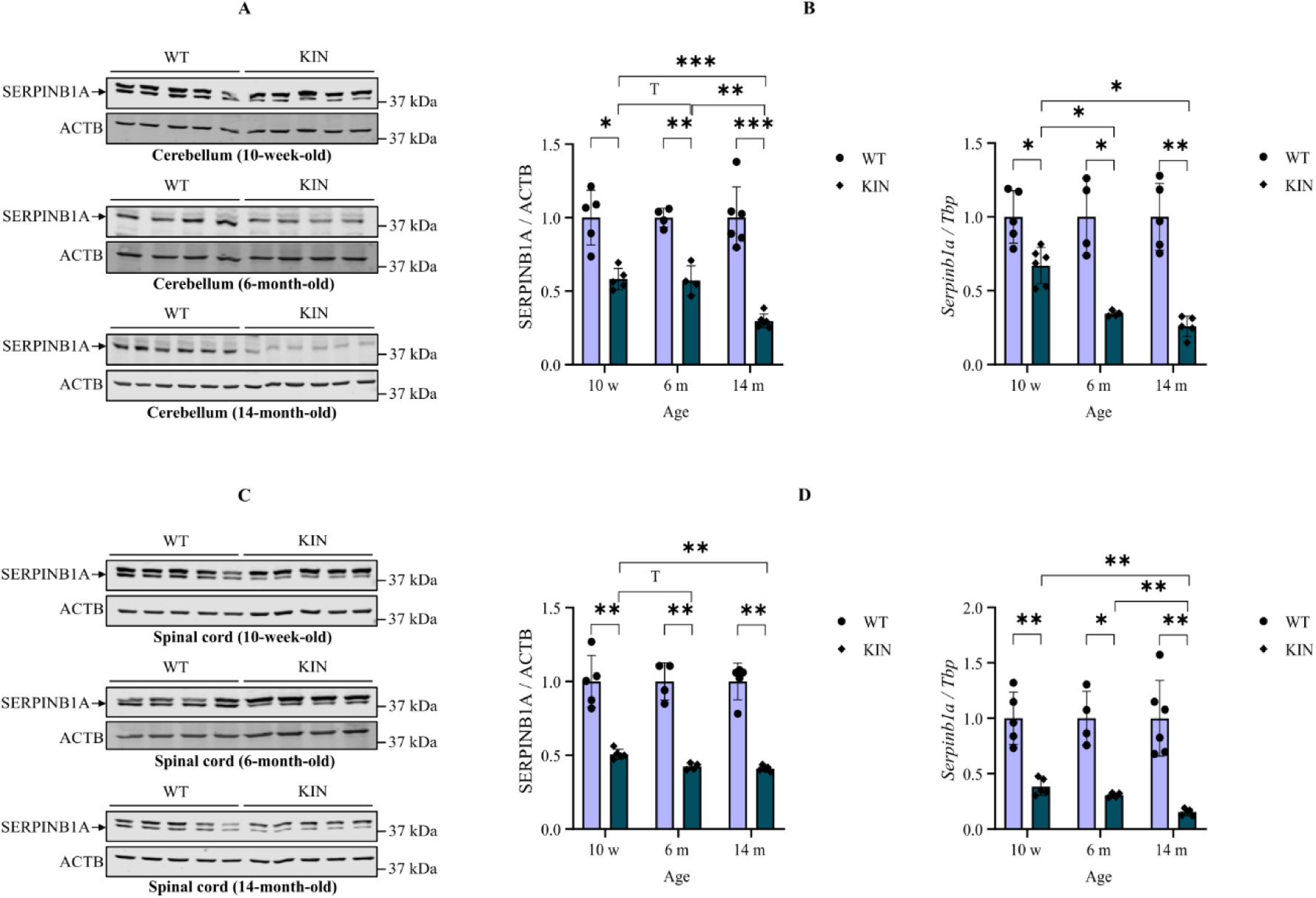
Quantitative immunoblots **(A, C)** and quantitative reverse-transcriptase real-time PCR **(B, D)** showing consistent downregulation of SERPINB1A/*Serpinb1a* in the cerebellum **(A, B)** and spinal cord **(C, D)** of 10-week-old (pre-manifest), six-month-old (early ataxic stage), and 14-month-old (late ataxic stage) *Atxn2*-CAG100-KIN mice. T < 0.10; *p < 0.05; ** p < 0.01, *** p < 0.001.

### SERPINB1 plasma levels are reduced and correlate with severity markers in SCA2 patients

Further validation efforts were conducted in 58 patients (male/female: 25/33) with clinical and molecular diagnosis of SCA2 and an equal number of healthy individuals from the same geographic area (male/female: 19/39). No significant differences were observed between SCA2 patients and controls regarding gender (χ^2^=1.318; p=0.251) and age distributions (t=- 0.688; p=0.493). Most patients were in disease stage 1 (mild ataxia) (62.07%), followed in frequency by patients with moderate ataxia (disease stage 2) (29.31%).

SERPINB1 blood plasma levels varied between 1.021 and 13.057 ng/ml in the studied sample, with a mean (SD) of 3.443 (2.107). Since SERPINB1 did not follow a normal distribution (K-S=0.140; p<0.001), it was log-transformed for hypothesis testing, then rendering an approximately normal distribution (K-S=0.077; p=0.097). Based on this, no significant differences were found for SERPINB1 plasma levels across genders (t=-0.091; p=0.927), and it was not significantly correlated with age (r=0.146; p=0.117). As a major finding, a nominally significant 17.88% reduction in SERPINB1 plasma levels was obtained for SCA2 patients relative to control individuals (t=2.618; p=0.010), with mean (SD) values of 3.070 (1.342) *versus* 4.051 (2.459) for patients and controls, respectively (Figure 2, A).

**Figure 2.**
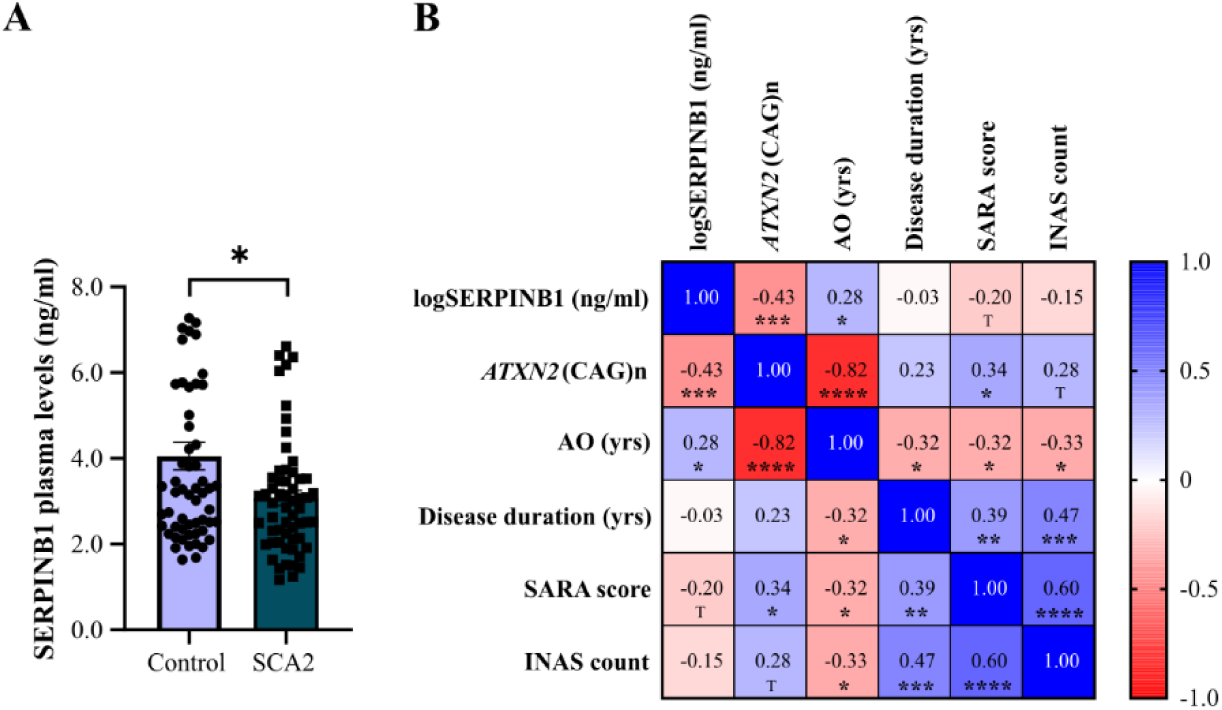
SERPINB1 plasma levels in SCA2 patients and healthy control individuals. **(A)** Mean comparisons for the log-transformed SERPINB1 plasma levels between patients (n=58) and age and gender matched (n=58) control individuals. **(B)** Correlation matrix for the log-transformed SERPINB1 plasma levels and disease severity markers in SCA2 patients (n=82). *p < 0.05; ** p < 0.01, *** p < 0.001, **** p < 0.0001; T− trend (0.10 > p > 0.05).

The relevance of SERPINB1 plasma levels for disease severity was assessed in an extended sample of 82 patients with SCA2 by linear correlation and regression analyses. Baseline characteristics for this cohort are shown in Table S6. Similar to the case-control cohort, most patients in the correlational study were in disease stage 1 (72.86%), followed in frequency by patients in disease stage 2 (17.14%). As expected, significant correlations were found between the CAG repeat length at expanded *ATXN2* alleles and clinical severity markers, including the age at onset and the SARA score. Remarkably, SERPINB1 plasma levels were significantly correlated with the CAG repeat length and the age at onset, and a trend was observed for the correlation between SERPINB1 plasma levels and SARA score (Figure 2, B). However, the influence of SERPINB1 plasma levels on the age at onset was no longer significant after adjusting it to the CAG repeat length by linear regression (t=0.061; p=0.952). Besides, a trend was observed for decreasing SERPINB1 plasma levels along with advancing disease stages, from mild (mean: 0.507 ng/ml), to moderate (mean: 0.437 ng/ml), to severe ataxia (mean: 0.251 ng/ml), though not statistically significant (F=1.544; p=0.221).

## Discussion

The search for reliable and informative biomarkers is crucial for assessing the clinical utility of experimental disease-modifying treatments for SCA2. Current clinical trials use ataxia scores, cerebellar imaging, nerve conductance, and perhaps the quantitation of NfL as axonal damage marker and of ATXN2 to assess efficacy of disease protein elimination, but molecular biomarkers in the affected pathways are lacking. Recent transcriptome profiling in nervous tissues from SCA2 mice models has revealed the crucial pathomechanisms (25, 69-72). As the novel contribution of the present study, we now explored the temporal evolution from pre-manifest to end-stage *Atxn2*-CAG100-KnockIn mice for transcripts involved in these processes, to define mechanistic disease progression biomarkers. Integrating multi-dimensional transcriptomic, proteomic, and phosphoproteomic profiles from cerebellar and spinal cord tissue, we looked for consistency across datasets. As the main finding, we define SERPINB1 downregulation in transcript, protein and phosphorylation assays as a promising candidate progression biomarker for SCA2.

Changes in *Serpinb1a* transcript levels and SERPINB1A protein abundance were verified by RT-qPCR and immunoblots, confirming their gradual decrease along with disease progression. Furthermore, using a sandwich enzyme-linked immunosorbent assay as an additional technique, a significant reduction in blood plasma SERPINB1 abundance was observed in SCA2 patients. Remarkably, blood plasma SERPINB1 abundance was significantly associated with the CAG repeat length at expanded *ATXN2* alleles and the age at disease onset, so lower SERPINB1 levels correlate with longer repeats and earlier onset as an indication of increased disease severity. Besides, a correlation of marginal significance with the SARA score reinforces the correlation of plasma SERPINB1 levels with clinical measures, although further studies in larger cohorts from different SCA2 populations will be needed.

SERPINB1 belongs to the clade B or ov-serpin subfamily of the SERPIN (serine or cysteine protease inhibitor) superfamily (73). It is needed to limit the excessive tissue damage during inflammatory responses to microbial infections by repressing the activity of elastase-like and chymotrypsin-like serine proteases and restricting the activity of inflammatory caspases (74-76). Besides, SERPINB1 may influence lysosomal pathways during immune responses (77). Although the functions of SERPINB1 in the nervous system have not been extensively studied, *SERPINB1* transcripts were found dysregulated in sporadic Creutzfeldt-Jakob disease and Alzheimer’s disease (AD) patients (78). According to genome-wide association studies of AD risk, genetic variants of SERPINB1 act as progression modifiers, and higher transcript levels of SERPINB1 in prefrontal cortex show direct correlation with increased amyloid Abeta 42 in cerebrospinal fluid (79). Furthermore, SERPINB1 was identified as a novel therapeutic target in a recent study using Mendelian randomization to repurpose licensed drugs for AD (80). These observations indicate that SERPINB1 exerts roles that are relevant for nervous system physiology and that its alteration is associated with neurodegenerative processes.

Although its overall expression levels in healthy brain are relatively low compared to other tissues, SERPINB1 is enriched in microglia, which is consistent with its role in inhibiting protease activity during inflammatory responses. Based on these observations, the progressive decreased abundance of SERPINB1 in the cerebellum and spinal cord of *Atxn2*-CAG100-KnockIn mice may indicate a state of dysregulated proteostasis, with de-repression of phagocytosis and lysosomal pathways in microglia as part of an innate immune response to neurons−derived polyQ-expanded Ataxin-2 aggregates, intended to limit cell-to-cell transfer of the aggregates and the spreading of pathology.

In line with this hypothesis, transcriptome profiling produced evidence of profuse *Serpins* downregulations along with lysosomal cathepsin upregulations in the cerebellum and spinal cord of *Atxn2*-CAG100-KnockIn mice. Given that the serpin superfamily of protease inhibitors primarily evolved as a buffering system to avoid excessive protein degradation by cathepsins and kallikrein-related peptidases (75, 81-84), these results suggest a simultaneous induction of cathepsins and suppression of serpins expression taking place. Remarkably, the transcriptional adaptation is significant and strong already at pre-manifest stages of our SCA2 mouse model, long before the ATXN2 aggregates become microscopically detectable in spinal cord and cerebellar neurons. Thus, the progressive lowering of SERPINB1 might become a clinical useful surrogate marker of microglial activation to prevent the formation of neuronal inclusion bodies. Similar compensatory cellular efforts may be a mechanism common to neurodegenerative proteinopathies, since dysregulations of diverse serpins and cathepsins were implicated in the pathology of several neurodegenerative disorders, including AD, PD, HD, ALS, and prion diseases (78, 85-90).

Interestingly, *Serpini1* transcript (encoding neuroserpin) showed consistent downregulations in the cerebellar and spinal cord KnockIn mice transcriptomes since early age, although to a lesser extent than *Serpinb1a*. Given the association of neuroserpin with neurodegenerative conditions (91, 92), and its involvement in the modulation of neurogenesis, neuronal axon growth and density of dendritic spines at synapses (93-96), it may become another relevant candidate biomarker for SCA2.

On the other hand, transcripts encoding cathepsins D, B, and L were among the most prominently upregulated in the nervous tissue of KnockIn mice. These cathepsins are the most abundant lysosomal proteases, ubiquitously expressed, and enriched in microglia (97, 98). Remarkably, these cathepsins, in addition to cathepsin Z, were implicated in the partial or total degradation of polyQ-expanded huntingtin (99-104). Interestingly, the *Ctsl* transcript showed the strongest progressive upregulations among all factors examined in the *Atxn2*-CAG100-KnockIn mice transcriptomes. Based on these observations, lysosomal cathepsins might play a role in the N-terminal proteolysis (105) or total degradation of polyQ-expanded Ataxin-2 monomers or aggregates, with potential relevance as SCA2 biomarkers and therapeutic targets. Besides cathepsin proteases, kallikrein-related peptidases (KLK) as another family of cellular proteases under control of serpins may play a significant role in SCA2 pathology. In particular, the *Klk6* transcript was progressively downregulated since early age. Validation studies provided additional evidence supporting a prominent reduced abundance of KLK6 protein in the cerebellum and spinal cord of end-stage KnockIn mice. Given that KLK6 promotes the degradation of myelin basic protein and plays a role in demyelinating conditions (106-108), apart from targeting protease−activated receptors PAR-1 and PAR-2 in response to neural damage (109), its downregulation in KnockIn mice seems to be protective. Similar to its proposed usefulness as a biomarker for AD (110-112), it is conceivable that KLK6 levels will be found to correlate with the widespread white matter damage in SCA2.

Further evidence for lysosome activation in *Atxn2*-CAG100-KnockIn mice due to polyQ-triggered protein aggregation toxicity comes from the observed strong upregulation of the microglia−enriched factors *Grn, Trem2*, *Laptm5*, and *Nfe2l2*, along with the prominent downregulation of *Itpr1*, *Arsg*, *Lamp5*, and *Kif5b* (113-120).

Conversely, significant upregulations of several translation factors involved in ribosome biogenesis and protein translation are reminiscent of observations in the Ataxin-2 Knock-Out mouse, thus suggesting cellular compensatory efforts in the face of an Ataxin-2 partial loss-of-function due to its mutation. Of particular interest among translation factors was the progressive induction of *Eif2ak2* in the spinal cord of KnockIn mice. *Eif2ak2* codes for the Eukaryotic Translation Initiation Factor 2 Alpha Kinase 2 (aka Protein Kinase R, PKR), which is a serine/threonine-specific, interferon-inducible protein kinase activated by viral or endogenous double-stranded RNA (dsRNA) and a crucial component of stress granules and the Integrated Stress Response (ISR) (121-123). In this scenario, the induction of *Eif2ak2* in the spinal cord of our *Atxn2*-CAG100-KnockIn mouse model may reflect a cellular innate immune system response elicited by double-stranded hairpin structures in the CAG-repeat expanded *Atxn2* mRNA, as a toxic RNA-mediated gain-of-function mechanism with a persistent downstream ISR dysregulation leading to impaired synaptic function and neural atrophy.

Altogether, in this study of temporal evolution in an unbiased multi-omic approach, with further validation studies, SERPINB1 was identified as novel promising biomarker with specificity for SCA2 pathomechanisms, whose decreased levels correlate with polyQ expansion size, age at onset, and show progressive reduction over time. Our findings suggest an early induction of microglial degradation efforts that result from mutant Ataxin-2 RNA toxicity and protein aggregation, involving the downregulation of protease inhibitors along with the upregulation of cathepsin proteases involved in the autophagy-lysosome pathway, which underlie SCA2 pathophysiology.

## Author Contributions

Conceptualization, S.G. and G.A.; methodology, J.C.-P., N.-E.S., L.- E.A.-M., and R.A.-R; validation, L.-E.A.-M.; formal analysis, G.A.; investigation, L.-E.A.-M., J.K., J.C.-P., N.-E.S., D.C., D.M., S.G.-S., D.C.-A., D.A.-G., L.M.O.-G., and R.A.-R.; resources, N.-E.S., J.C.-P., S.G., and J.M.-M.; data curation, G.A.; writing—original draft preparation, L.E.A.-M., and G.A.; writing—review and editing, N.-E.S., J.K., and S.G.; visualization, L.-E.A.-M., and G.A.; project administration, G.A.; funding acquisition, G.A. All authors have read and agreed to the published version of the manuscript.

## Financial Disclosures

None.

## Supplementary

**Figure S1.** Volcano plots showing transcriptionally dysregulated translation-related factors in the cerebellum **(A, B)** and spinal cord **(C, D)** of pre-manifest (10-week-old) **(A, C)** or end-stage (14-month-old) **(B, D)** *Atxn2*-CAG100-KnockIn mice. Down- and upregulated transcripts with at least 1.2-fold expression levels changes in each direction reaching nominal significance in each dataset were colored in green and red, respectively. Among upregulated transcripts, only representative factors with more prominent dysregulations were labelled. Factors with higher than 2.0-fold dysregulations in each direction are shown in bold.

**Figure S2.** Volcano plots showing transcriptionally dysregulated stress granule and RNA processing factors in the cerebellum (A, B) and spinal cord (C, D) of pre-manifest (10-week-old) (A, C) or end-stage (14-month-old) (B, D) *Atxn2*-CAG100-KnockIn mice. Down- and upregulated transcripts with at least 1.2-fold expression levels changes in each direction reaching nominal significance in each dataset were colored in green and red, respectively. Only factors with more prominent dysregulations (FC 1.5 in each direction) were labelled. Factors with higher than 2.0-fold dysregulations in each direction are shown in bold.

**Figure S3.** Volcano plots showing transcriptionally dysregulated factors of the Autophagy-Lysosome pathway in the cerebellum **(A, B)** and spinal cord **(C, D)** of pre-manifest (10-week-old) **(A, C)** or end-stage (14-month-old) **(B, D)** *Atxn2*-CAG100-KnockIn mice. Down- and upregulated transcripts with at least 1.2-fold expression levels changes in each direction reaching nominal significance in each dataset were colored in green and red, respectively. Only factors with more prominent dysregulations (FC 1.5 in each direction) were labelled. Factors with higher than 2.0-fold dysregulations in each direction are shown in bold, and those with higher than 3.0-fold dysregulations were underlined.

**Figure S4.** Quantitative reverse-transcriptase real-time PCR (N = 6 vs. 6) for transcripts encoding cathepsin proteinases as the main mammalian lysosomal proteases, in end-stage cerebellum **(A)** and spinal cord **(B)** *Atxn2*-CAG100-KnockIn mice. * p < 0.05; ** p < 0.01, *** p < 0.001, **** p < 0.0001, ns = not significant.

**Figure S5.** Quantitative immunoblots **(A, C)** and quantitative reverse-transcriptase real-time PCR **(B, D)** showing consistent downregulation of KLK6/*Klk6* in the cerebellum **(A, B)** and spinal cord **(C, D)** of 10-week-old (pre-manifest), six-month-old (early ataxic stage), and 14-month-old (late ataxic stage) *Atxn2*-CAG100-KIN mice. w = week, m = month. T < 0.10; *p < 0.05; ** p < 0.01, *** p < 0.001.

**Figure S6.** Commonly dysregulated factors in pre-manifest and end-stage *Atxn2*-CAG100-KnockIn mice, based on FC ≥1.5 in each direction, and nominal significance (p<0.05). Venn diagrams of the up- **(A)** and down-regulated **(B)** transcripts in the cerebellar (T_Cbll) and spinal cord transcriptomes (T_SC), and cerebellar proteome (P_Cbll) of pre-manifest KnockIn mice. Venn diagrams of the up- **(C)** or down-regulated **(D)** transcripts/proteins in the cerebellar transcriptome (T_Cbll) and proteome (P_Cbll), and spinal cord transcriptome (T_SC), proteome (P_SC), and spinal cord phosphoproteome (PP_SC) of end-stage KnockIn mice.

**Figure S7.** Volcano plots showing transcriptionally dysregulated Serpins in the cerebellum **(A, B)** and spinal cord **(C, D)** of pre-manifest (10-week-old) **(A, C)** or end-stage (14-month-old) **(B, D)** *Atxn2*-CAG100-KnockIn mice. Down- and upregulated transcripts reaching nominal significance in each dataset were labeled and colored in green and red, respectively. Dashed lines represent the cutoff p-value for nominal significance (-log10 p-value = 1.3).

**Figure S8.** Quantitative reverse-transcriptase real-time PCR (N = 5 vs. 5) for transcripts encoding serpin protease inhibitors, in the cerebellum **(A, C, E)** and spinal cord **(B, D, F)** of *Atxn2*-CAG100-KnockIn mice. Experiments in **A** and **B** were conducted in end-stage mice (14-month-old), while experiments in **C-F** were conducted in pre-manifest (10-week-old) and end-stage mice (14-month-old) mice. * p < 0.05; ** p < 0.01, *** p < 0.001, **** p < 0.0001, ns = not significant.

**Table S1.** List of transcriptionally dysregulated translation-related factors in the cerebellum and spinal cord of pre-manifest (10-week-old) or end-stage (14-month-old) *Atxn2*-CAG100-KnockIn mice. Down- and upregulated transcripts with at least 1.2-fold expression levels changes in each direction reaching nominal significance in each dataset are shown.

**Table S2.** List of transcriptionally dysregulated translation, stress granules, RNA processing, and autophago-lysosome pathway factors progressing over *Atxn2*-CAG100-KnockIn lifespan. Candidate progression markers were selected from global transcriptome profiles upon ≥1.5-fold change at 14 months of age, preceded by also significant >1.2-fold expression change at 10 weeks of age, in *Atxn2*-CAG100-KIN cerebellar and cervicothoracic spinal cord tissue.

**Table S3.** List of transcriptionally dysregulated stress granules and RNA processing factors in the cerebellum and spinal cord of pre-manifest (10-week-old) or end-stage (14-month-old) *Atxn2*-CAG100-KnockIn mice. Down- and upregulated transcripts with at least 1.2-fold expression levels changes in each direction reaching nominal significance in each dataset are shown.

**Table S4.** List of transcriptionally dysregulated autophago-lysosome factors in the cerebellum and spinal cord of pre-manifest (10-week-old) or end-stage (14-month-old) *Atxn2*-CAG100-KnockIn mice. Down- and upregulated transcripts with at least 1.2-fold expression levels changes in each direction reaching nominal significance in each dataset are shown.

**Table S5.** List of transcriptionally dysregulated serpins in the cerebellum and spinal cord of pre-manifest (10-week-old) or end-stage (14-month-old) *Atxn2*-CAG100-KnockIn mice. Down- and upregulated transcripts with at least 1.2-fold expression levels changes in each direction reaching nominal significance in each dataset are shown.

**Table S6.** Clinical and molecular characteristics of the extended cohort of patients with SCA2.

## Notes

**Conflict of Interest:** The authors declare no conflicts of interest. The funders had no role in the design of the study; in the collection, analyses, or interpretation of data; in the writing of the manuscript; or in the decision to publish the results.

### Competing Interest Statement

The authors have declared no competing interest.

## References

1. Imbert G, Saudou F, Yvert G, Devys D, Trottier Y, Garnier JM, et al. Cloning of the gene for spinocerebellar ataxia 2 reveals a locus with high sensitivity to expanded CAG/glutamine repeats. Nat Genet. 1996;14(3):285–91.

2. Pulst SM, Nechiporuk A, Nechiporuk T, Gispert S, Chen XN, Lopes-Cendes I, et al. Moderate expansion of a normally biallelic trinucleotide repeat in spinocerebellar ataxia type 2. Nat Genet. 1996;14(3):269–76.

3. Sanpei K, Takano H, Igarashi S, Sato T, Oyake M, Sasaki H, et al. Identification of the spinocerebellar ataxia type 2 gene using a direct identification of repeat expansion and cloning technique, DIRECT. Nat Genet. 1996;14(3):277–84.

4. Almaguer-Mederos LE, Pérez-Ávila I, Aguilera-Rodríguez R, Velázquez-Garcés M, Almaguer-Gotay D, Hechavarría-Pupo R, et al. Body Mass Index Is Significantly Associated With Disease Severity in Spinocerebellar Ataxia Type 2 Patients. Mov Disord. 2021;36(6):1372–80.

5. Almaguer-Mederos LE, Aguilera Rodríguez R, González Zaldivar Y, Almaguer Gotay D, Cuello Almarales D, Laffita Mesa J, et al. Estimation of survival in spinocerebellar ataxia type 2 Cuban patients. Clin Genet. 2013;83(3):293–4.

6. Orozco Diaz G, Nodarse Fleites A, Cordovés Sagaz R, Auburger G. Autosomal dominant cerebellar ataxia: clinical analysis of 263 patients from a homogeneous population in Holguín, Cuba. Neurology. 1990;40(9):1369–75.

7. Rüb U, Schöls L, Paulson H, Auburger G, Kermer P, Jen JC, et al. Clinical features, neurogenetics and neuropathology of the polyglutamine spinocerebellar ataxias type 1, 2, 3, 6 and 7. Prog Neurobiol. 2013;104:38-66.

8. Yang L, Dong Y, Ma Y, Ni W, Wu ZY. Genetic profile and clinical characteristics of Chinese patients with spinocerebellar ataxia type 2: A multicenter experience over 10 years. Eur J Neurol. 2021;28(3):955–64.

9. Elden AC, Kim HJ, Hart MP, Chen-Plotkin AS, Johnson BS, Fang X, et al. Ataxin-2 intermediate-length polyglutamine expansions are associated with increased risk for ALS. Nature. 2010;466(7310):1069-75.

10. Glass JD, Dewan R, Ding J, Gibbs JR, Dalgard C, Keagle PJ, et al. ATXN2 intermediate expansions in amyotrophic lateral sclerosis. Brain. 2022;145(8):2671–6.

11. Chiò A, Moglia C, Canosa A, Manera U, Grassano M, Vasta R, et al. Association of Copresence of Pathogenic Variants Related to Amyotrophic Lateral Sclerosis and Prognosis. Neurology. 2023;101(1):e83–e93.

12. Rubino E, Mancini C, Boschi S, Ferrero P, Ferrone M, Bianca S, et al. ATXN2 intermediate repeat expansions influence the clinical phenotype in frontotemporal dementia. Neurobiol Aging. 2019;73:231.e7-.e9.

13. Fournier C, Anquetil V, Camuzat A, Stirati-Buron S, Sazdovitch V, Molina-Porcel L, et al. Interrupted CAG expansions in ATXN2 gene expand the genetic spectrum of frontotemporal dementias. Acta Neuropathol Commun. 2018;6(1):41.

14. Ross OA, Rutherford NJ, Baker M, Soto-Ortolaza AI, Carrasquillo MM, DeJesus-Hernandez M, et al. Ataxin-2 repeat-length variation and neurodegeneration. Hum Mol Genet. 2011;20(16):3207–12.

15. Kim YE, Jeon B, Farrer MJ, Scott E, Guella I, Park SS, et al. SCA2 family presenting as typical Parkinson’s disease: 34 year follow up. Parkinsonism Relat Disord. 2017;40:69–72.

16. Almaguer-Mederos L-E, Kandi AR, Sen N-E, Canet-Pons J, Berger L-M, Key J, et al. Spinal cord phosphoproteome of a SCA2/ALS13 mouse model reveals alteration of ATXN2-N-term SH3-actin interactome and of autophagy via WNK1-MYO6-OPTN-SQSTM1. bioRxiv. 2024:2024.11.06.622233.

17. Del Castillo U, Norkett R, Lu W, Serpinskaya A, Gelfand VI. Ataxin-2 is essential for cytoskeletal dynamics and neurodevelopment in Drosophila. iScience. 2022;25(1):103536.

18. Inagaki H, Hosoda N, Tsuiji H, Hoshino SI. Direct evidence that Ataxin-2 is a translational activator mediating cytoplasmic polyadenylation. J Biol Chem. 2020;295(47):15810–25.

19. Lastres-Becker I, Nonis D, Eich F, Klinkenberg M, Gorospe M, Kötter P, et al. Mammalian ataxin-2 modulates translation control at the pre-initiation complex via PI3K/mTOR and is induced by starvation. Biochim Biophys Acta. 2016;1862(9):1558–69.

20. Liu YJ, Wang JY, Zhang XL, Jiang LL, Hu HY. Ataxin-2 sequesters Raptor into aggregates and impairs cellular mTORC1 signaling. Febs j. 2024;291(8):1795–812.

21. Nonis D, Schmidt MHH, van de Loo S, Eich F, Dikic I, Nowock J, Auburger G. Ataxin-2 associates with the endocytosis complex and affects EGF receptor trafficking. Cell Signal. 2008;20(10):1725–39.

22. Satterfield TF, Jackson SM, Pallanck LJ. A Drosophila homolog of the polyglutamine disease gene SCA2 is a dosage-sensitive regulator of actin filament formation. Genetics. 2002;162(4):1687–702.

23. Yamagishi R, Inagaki H, Suzuki J, Hosoda N, Sugiyama H, Tomita K, et al. Concerted action of ataxin-2 and PABPC1-bound mRNA poly(A) tail in the formation of stress granules. Nucleic Acids Res. 2024;52(15):9193–209.

24. Rüb U, Del Turco D, Del Tredici K, de Vos RA, Brunt ER, Reifenberger G, et al. Thalamic involvement in a spinocerebellar ataxia type 2 (SCA2) and a spinocerebellar ataxia type 3 (SCA3) patient, and its clinical relevance. Brain. 2003;126(Pt 10):2257–72.

25. Sen NE, Canet-Pons J, Halbach MV, Arsovic A, Pilatus U, Chae WH, et al. Generation of an Atxn2-CAG100 knock-in mouse reveals N-acetylaspartate production deficit due to early Nat8l dysregulation. Neurobiol Dis. 2019;132:104559.

26. Scoles DR, Gandelman M, Paul S, Dexheimer T, Dansithong W, Figueroa KP, et al. A quantitative high-throughput screen identifies compounds that lower expression of the SCA2-and ALS-associated gene ATXN2. J Biol Chem. 2022;298(8):102228.

27. Rodriguez CM, Bechek SC, Jones GL, Nakayama L, Akiyama T, Kim G, et al. Targeting RTN4/NoGo-Receptor reduces levels of ALS protein ataxin-2. Cell Rep. 2022;41(4):111505.

28. Kim G, Nakayama L, Blum JA, Akiyama T, Boeynaems S, Chakraborty M, et al. Genome-wide CRISPR screen reveals v-ATPase as a drug target to lower levels of ALS protein ataxin-2. Cell Rep. 2022;41(4):111508.

29. Scoles DR, Meera P, Schneider MD, Paul S, Dansithong W, Figueroa KP, et al. Antisense oligonucleotide therapy for spinocerebellar ataxia type 2. Nature. 2017;544(7650):362-6.

30. Becker LA, Huang B, Bieri G, Ma R, Knowles DA, Jafar-Nejad P, et al. Therapeutic reduction of ataxin-2 extends lifespan and reduces pathology in TDP-43 mice. Nature. 2017;544(7650):367-71.

31. Zeballos CM, Moore HJ, Smith TJ, Powell JE, Ahsan NS, Zhang S, Gaj T. Mitigating a TDP-43 proteinopathy by targeting ataxin-2 using RNA-targeting CRISPR effector proteins. Nat Commun. 2023;14(1):6492.

32. Wardman JH, Henriksen EE, Marthaler AG, Nielsen JE, Nielsen TT. Enhancement of Autophagy and Solubilization of Ataxin-2 Alleviate Apoptosis in Spinocerebellar Ataxia Type 2 Patient Cells. Cerebellum. 2020;19(2):165–81.

33. Southwell AL, Smith SE, Davis TR, Caron NS, Villanueva EB, Xie Y, et al. Ultrasensitive measurement of huntingtin protein in cerebrospinal fluid demonstrates increase with Huntington disease stage and decrease following brain huntingtin suppression. Sci Rep. 2015;5:12166.

34. Fischer DF, Dijkstra S, Lo K, Suijker J, Correia ACP, Naud P, et al. Development of mAb-based polyglutamine-dependent and polyglutamine length-independent huntingtin quantification assays with cross-site validation. PLoS One. 2022;17(4):e0266812.

35. Hübener-Schmid J, Kuhlbrodt K, Peladan J, Faber J, Santana MM, Hengel H, et al. Polyglutamine-Expanded Ataxin-3: A Target Engagement Marker for Spinocerebellar Ataxia Type 3 in Peripheral Blood. Mov Disord. 2021;36(11):2675–81.

36. The Genotype-Tissue Expression (GTEx) project. Nat Genet. 2013;45(6):580-5.

37. Bux J, Sen NE, Klink IM, Hauser S, Synofzik M, Schöls L, et al. TR-FRET-Based Immunoassay to Measure Ataxin-2 as a Target Engagement Marker in Spinocerebellar Ataxia Type 2. Mol Neurobiol. 2023;60(6):3553–67.

38. Sen NE, Drost J, Gispert S, Torres-Odio S, Damrath E, Klinkenberg M, et al. Search for SCA2 blood RNA biomarkers highlights Ataxin-2 as strong modifier of the mitochondrial factor PINK1 levels. Neurobiol Dis. 2016;96:115–26.

39. Puorro G, Marsili A, Sapone F, Pane C, De Rosa A, Peluso S, et al. Peripheral markers of autophagy in polyglutamine diseases. Neurol Sci. 2018;39(1):149–52.

40. Squadrone S, Brizio P, Mancini C, Abete MC, Brusco A. Altered homeostasis of trace elements in the blood of SCA2 patients. J Trace Elem Med Biol. 2018;47:111–4.

41. Dennis AG, Almaguer-Mederos LE, Raúl RA, Roberto RL, Luis VP, Dany CA, et al. Redox Imbalance Associates with Clinical Worsening in Spinocerebellar Ataxia Type 2. Oxid Med Cell Longev. 2021;2021:9875639.

42. Almaguer-Mederos LE, Aguilera-Rodríguez R, Almaguer-Gotay D, Hechavarría-Barzaga K, Álvarez-Sosa A, Chapman-Rodríguez Y, et al. Testosterone Levels Are Decreased and Associated with Disease Duration in Male Spinocerebellar Ataxia Type 2 Patients. Cerebellum. 2020;19(4):597–604.

43. Shen XN, Wu KM, Huang YY, Guo Y, Huang SY, Zhang YR, et al. Systematic assessment of plasma biomarkers in spinocerebellar ataxia. Neurobiol Dis. 2023;181:106112.

44. Yang L, Shao YR, Li XY, Ma Y, Dong Y, Wu ZY. Association of the Level of Neurofilament Light With Disease Severity in Patients With Spinocerebellar Ataxia Type 2. Neurology. 2021;97(24):e2404–e13.

45. Coarelli G, Darios F, Petit E, Dorgham K, Adanyeguh I, Petit E, et al. Plasma neurofilament light chain predicts cerebellar atrophy and clinical progression in spinocerebellar ataxia. Neurobiol Dis. 2021;153:105311.

46. Coarelli G, Dubec-Fleury C, Petit E, Sayah S, Fischer C, Nassisi M, et al. Longitudinal Changes of Clinical, Imaging, and Fluid Biomarkers in Preataxic and Early Ataxic Spinocerebellar Ataxia Type 2 and 7 Carriers. Neurology. 2024;103(5):e209749.

47. Peng L, Wang S, Chen Z, Peng Y, Wang C, Long Z, et al. Blood Neurofilament Light Chain in Genetic Ataxia: A Meta-Analysis. Mov Disord. 2022;37(1):171–81.

48. Sun Q, Zhao X, Li S, Yang F, Wang H, Cui F, Huang X. CSF Neurofilament Light Chain Elevation Predicts ALS Severity and Progression. Front Neurol. 2020;11:919.

49. Verde F, Milone I, Colombo E, Maranzano A, Solca F, Torre S, et al. Phenotypic correlates of serum neurofilament light chain levels in amyotrophic lateral sclerosis. Front Aging Neurosci. 2023;15:1132808.

50. Machacek M, Garcia-Montoya E, McColgan P, Sanwald-Ducray P, Mazer NA. NfL concentration in CSF is a quantitative marker of the rate of neurodegeneration in aging and Huntington’s disease: a semi-mechanistic model-based analysis. Front Neurosci. 2024;18:1420198.

51. Tezenas du Montcel S, Petit E, Olubajo T, Faber J, Lallemant-Dudek P, Bushara K, et al. Baseline Clinical and Blood Biomarkers in Patients With Preataxic and Early-Stage Disease Spinocerebellar Ataxia 1 and 3. Neurology. 2023;100(17):e1836–e48.

52. Miller TM, Cudkowicz ME, Genge A, Shaw PJ, Sobue G, Bucelli RC, et al. Trial of Antisense Oligonucleotide Tofersen for SOD1 ALS. N Engl J Med. 2022;387(12):1099–110.

53. Meyer T, Schumann P, Weydt P, Petri S, Koc Y, Spittel S, et al. Neurofilament light-chain response during therapy with antisense oligonucleotide tofersen in SOD1-related ALS: Treatment experience in clinical practice. Muscle Nerve. 2023;67(6):515–21.

54. McColgan P, Thobhani A, Boak L, Schobel SA, Nicotra A, Palermo G, et al. Tominersen in Adults with Manifest Huntington’s Disease. N Engl J Med. 2023;389(23):2203–5.

55. Tabrizi SJ, Leavitt BR, Landwehrmeyer GB, Wild EJ, Saft C, Barker RA, et al. Targeting Huntingtin Expression in Patients with Huntington’s Disease. N Engl J Med. 2019;380(24):2307–16.

56. Carraro C, Montgomery JV, Klimmt J, Paquet D, Schultze JL, Beyer MD. Tackling neurodegeneration in vitro with omics: a path towards new targets and drugs. Front Mol Neurosci. 2024;17:1414886.

57. Di Paolo A, Garat J, Eastman G, Farias J, Dajas-Bailador F, Smircich P, Sotelo-Silveira JR. Functional Genomics of Axons and Synapses to Understand Neurodegenerative Diseases. Front Cell Neurosci. 2021;15:686722.

58. Meng L, Jin H, Yulug B, Altay O, Li X, Hanoglu L, et al. Multi-omics analysis reveals the key factors involved in the severity of the Alzheimer’s disease. Alzheimers Res Ther. 2024;16(1):213.

59. Carrillo F, Palomba NP, Ghirimoldi M, Didò C, Fortunato G, Khoso S, et al. Multiomics approach discloses lipids and metabolites profiles associated to Parkinson’s disease stages and applied therapies. Neurobiol Dis. 2024;202:106698.

60. Mitropoulos K, Katsila T, Patrinos GP, Pampalakis G. Multi-Omics for Biomarker Discovery and Target Validation in Biofluids for Amyotrophic Lateral Sclerosis Diagnosis. Omics. 2018;22(1):52–64.

61. Paryani F, Kwon JS, Ng CW, Jakubiak K, Madden N, Ofori K, et al. Multi-omic analysis of Huntington’s disease reveals a compensatory astrocyte state. Nat Commun. 2024;15(1):6742.

62. Fittschen M, Lastres-Becker I, Halbach MV, Damrath E, Gispert S, Azizov M, et al. Genetic ablation of ataxin-2 increases several global translation factors in their transcript abundance but decreases translation rate. Neurogenetics. 2015;16(3):181–92.

63. Zhuang Y, Li Z, Xiong S, Sun C, Li B, Wu SA, et al. Circadian clocks are modulated by compartmentalized oscillating translation. Cell. 2023;186(15):3245–60.e23.

64. Nonhoff U, Ralser M, Welzel F, Piccini I, Balzereit D, Yaspo ML, et al. Ataxin-2 interacts with the DEAD/H-box RNA helicase DDX6 and interferes with P-bodies and stress granules. Mol Biol Cell. 2007;18(4):1385–96.

65. Swisher KD, Parker R. Localization to, and effects of Pbp1, Pbp4, Lsm12, Dhh1, and Pab1 on stress granules in Saccharomyces cerevisiae. PLoS One. 2010;5(4):e10006.

66. Yokoshi M, Li Q, Yamamoto M, Okada H, Suzuki Y, Kawahara Y. Direct binding of Ataxin-2 to distinct elements in 3’ UTRs promotes mRNA stability and protein expression. Mol Cell. 2014;55(2):186–98.

67. Drost J, Nonis D, Eich F, Leske O, Damrath E, Brunt ER, et al. Ataxin-2 modulates the levels of Grb2 and SRC but not ras signaling. J Mol Neurosci. 2013;51(1):68–81.

68. Canet-Pons J, Schubert R, Duecker RP, Schrewe R, Wölke S, Kieslich M, et al. Ataxia telangiectasia alters the ApoB and reelin pathway. Neurogenetics. 2018;19(4):237–55.

69. Arsović A, Halbach MV, Canet-Pons J, Esen-Sehir D, Döring C, Freudenberg F, et al. Mouse Ataxin-2 Expansion Downregulates CamKII and Other Calcium Signaling Factors, Impairing Granule-Purkinje Neuron Synaptic Strength. Int J Mol Sci. 2020;21(18).

70. Canet-Pons J, Sen NE, Arsović A, Almaguer-Mederos LE, Halbach MV, Key J, et al. Atxn2-CAG100-KnockIn mouse spinal cord shows progressive TDP43 pathology associated with cholesterol biosynthesis suppression. Neurobiol Dis. 2021;152:105289.

71. Scoles DR, Dansithong W, Pflieger LT, Paul S, Gandelman M, Figueroa KP, et al. ALS-associated genes in SCA2 mouse spinal cord transcriptomes. Hum Mol Genet. 2020;29(10):1658–72.

72. Pflieger LT, Dansithong W, Paul S, Scoles DR, Figueroa KP, Meera P, et al. Gene co-expression network analysis for identifying modules and functionally enriched pathways in SCA2. Hum Mol Genet. 2017;26(16):3069–80.

73. Silverman GA, Whisstock JC, Askew DJ, Pak SC, Luke CJ, Cataltepe S, et al. Human clade B serpins (ov-serpins) belong to a cohort of evolutionarily dispersed intracellular proteinase inhibitor clades that protect cells from promiscuous proteolysis. Cell Mol Life Sci. 2004;61(3):301–25.

74. Choi YJ, Kim S, Choi Y, Nielsen TB, Yan J, Lu A, et al. SERPINB1-mediated checkpoint of inflammatory caspase activation. Nat Immunol. 2019;20(3):276–87.

75. Burgener SS, Leborgne NGF, Snipas SJ, Salvesen GS, Bird PI, Benarafa C. Cathepsin G Inhibition by Serpinb1 and Serpinb6 Prevents Programmed Necrosis in Neutrophils and Monocytes and Reduces GSDMD-Driven Inflammation. Cell Rep. 2019;27(12):3646–56.e5.

76. Baumann M, Pham CT, Benarafa C. SerpinB1 is critical for neutrophil survival through cell-autonomous inhibition of cathepsin G. Blood. 2013;121(19):3900–7, s1-6.

77. Yan J, Gao Y, Bai J, Li J, Li M, Liu X, Jiang P. SERPINB1 promotes Senecavirus A replication by degrading IKBKE and regulating the IFN pathway via autophagy. J Virol. 2023;97(10):e0104523.

78. Zattoni M, Mearelli M, Vanni S, Colini Baldeschi A, Tran TH, Ferracin C, et al. Serpin Signatures in Prion and Alzheimer’s Diseases. Mol Neurobiol. 2022;59(6):3778–99.

79. Deming Y, Dumitrescu L, Barnes LL, Thambisetty M, Kunkle B, Gifford KA, et al. Sex-specific genetic predictors of Alzheimer’s disease biomarkers. Acta Neuropathol. 2018;136(6):857–72.

80. Su WM, Gu XJ, Dou M, Duan QQ, Jiang Z, Yin KF, et al. Systematic druggable genome-wide Mendelian randomisation identifies therapeutic targets for Alzheimer’s disease. J Neurol Neurosurg Psychiatry. 2023;94(11):954–61.

81. Janciauskiene S, Lechowicz U, Pelc M, Olejnicka B, Chorostowska-Wynimko J. Diagnostic and therapeutic value of human serpin family proteins. Biomed Pharmacother. 2024;175:116618.

82. Masurier N, Arama DP, El Amri C, Lisowski V. Inhibitors of kallikrein-related peptidases: An overview. Med Res Rev. 2018;38(2):655–83.

83. Huntington JA, Read RJ, Carrell RW. Structure of a serpin-protease complex shows inhibition by deformation. Nature. 2000;407(6806):923-6.

84. Sánchez-Navarro A, González-Soria I, Caldiño-Bohn R, Bobadilla NA. An integrative view of serpins in health and disease: the contribution of SerpinA3. Am J Physiol Cell Physiol. 2021;320(1):C106–c18.

85. Hällqvist J, Bartl M, Dakna M, Schade S, Garagnani P, Bacalini MG, et al. Plasma proteomics identify biomarkers predicting Parkinson’s disease up to 7 years before symptom onset. Nat Commun. 2024;15(1):4759.

86. Onur TS, Laitman A, Zhao H, Keyho R, Kim H, Wang J, et al. Downregulation of glial genes involved in synaptic function mitigates Huntington’s disease pathogenesis. Elife. 2021;10.

87. Humphrey J, Venkatesh S, Hasan R, Herb JT, de Paiva Lopes K, Küçükali F, et al. Integrative transcriptomic analysis of the amyotrophic lateral sclerosis spinal cord implicates glial activation and suggests new risk genes. Nat Neurosci. 2023;26(1):150–62.

88. Cermak S, Kosicek M, Mladenovic-Djordjevic A, Smiljanic K, Kanazir S, Hecimovic S. Loss of Cathepsin B and L Leads to Lysosomal Dysfunction, NPC-Like Cholesterol Sequestration and Accumulation of the Key Alzheimer’s Proteins. PLoS One. 2016;11(11):e0167428.

89. Drobny A, Prieto Huarcaya S, Dobert J, Kluge A, Bunk J, Schlothauer T, Zunke F. The role of lysosomal cathepsins in neurodegeneration: Mechanistic insights, diagnostic potential and therapeutic approaches. Biochim Biophys Acta Mol Cell Res. 2022;1869(7):119243.

90. Prieto Huarcaya S, Drobny A, Marques ARA, Di Spiezio A, Dobert JP, Balta D, et al. Recombinant pro-CTSD (cathepsin D) enhances SNCA/α-Synuclein degradation in α-Synucleinopathy models. Autophagy. 2022;18(5):1127–51.

91. Eshima J, O’Connor SA, Marschall E, Bowser R, Plaisier CL, Smith BS. Molecular subtypes of ALS are associated with differences in patient prognosis. Nat Commun. 2023;14(1):95.

92. D’Acunto E, Fra A, Visentin C, Manno M, Ricagno S, Galliciotti G, Miranda E. Neuroserpin: structure, function, physiology and pathology. Cell Mol Life Sci. 2021;78(19-20):6409–30.

93. Parmar PK, Coates LC, Pearson JF, Hill RM, Birch NP. Neuroserpin regulates neurite outgrowth in nerve growth factor-treated PC12 cells. J Neurochem. 2002;82(6):1406–15.

94. Osterwalder T, Cinelli P, Baici A, Pennella A, Krueger SR, Schrimpf SP, et al. The axonally secreted serine proteinase inhibitor, neuroserpin, inhibits plasminogen activators and plasmin but not thrombin. J Biol Chem. 1998;273(4):2312–21.

95. Godinez A, Rajput R, Chitranshi N, Gupta V, Basavarajappa D, Sharma S, et al. Neuroserpin, a crucial regulator for axogenesis, synaptic modelling and cell-cell interactions in the pathophysiology of neurological disease. Cell Mol Life Sci. 2022;79(3):172.

96. Hermann M, Reumann R, Schostak K, Kement D, Gelderblom M, Bernreuther C, et al. Deficits in developmental neurogenesis and dendritic spine maturation in mice lacking the serine protease inhibitor neuroserpin. Mol Cell Neurosci. 2020;102:103420.

97. Nakanishi H. Cathepsin regulation on microglial function. Biochim Biophys Acta Proteins Proteom. 2020;1868(9):140465.

98. Yadati T, Houben T, Bitorina A, Shiri-Sverdlov R. The Ins and Outs of Cathepsins: Physiological Function and Role in Disease Management. Cells. 2020;9(7).

99. Kegel KB, Kim M, Sapp E, McIntyre C, Castaño JG, Aronin N, DiFiglia M. Huntingtin expression stimulates endosomal-lysosomal activity, endosome tubulation, and autophagy. J Neurosci. 2000;20(19):7268–78.

100. Kegel KB, Sapp E, Alexander J, Reeves P, Bleckmann D, Sobin L, et al. Huntingtin cleavage product A forms in neurons and is reduced by gamma-secretase inhibitors. Mol Neurodegener. 2010;5:58.

101. Kim YJ, Sapp E, Cuiffo BG, Sobin L, Yoder J, Kegel KB, et al. Lysosomal proteases are involved in generation of N-terminal huntingtin fragments. Neurobiol Dis. 2006;22(2):346–56.

102. Liang Q, Ouyang X, Schneider L, Zhang J. Reduction of mutant huntingtin accumulation and toxicity by lysosomal cathepsins D and B in neurons. Mol Neurodegener. 2011;6:37.

103. Qin ZH, Wang Y, Kegel KB, Kazantsev A, Apostol BL, Thompson LM, et al. Autophagy regulates the processing of amino terminal huntingtin fragments. Hum Mol Genet. 2003;12(24):3231–44.

104. Bhutani N, Piccirillo R, Hourez R, Venkatraman P, Goldberg AL. Cathepsins L and Z are critical in degrading polyglutamine-containing proteins within lysosomes. J Biol Chem. 2012;287(21):17471–82.

105. Chitre M, Emery P. ATXN2 is a target of N-terminal proteolysis. PLoS One. 2023;18(12):e0296085.

106. Bernett MJ, Blaber SI, Scarisbrick IA, Dhanarajan P, Thompson SM, Blaber M. Crystal structure and biochemical characterization of human kallikrein 6 reveals that a trypsin-like kallikrein is expressed in the central nervous system. J Biol Chem. 2002;277(27):24562–70.

107. Blaber SI, Scarisbrick IA, Bernett MJ, Dhanarajan P, Seavy MA, Jin Y, et al. Enzymatic properties of rat myelencephalon-specific protease. Biochemistry. 2002;41(4):1165–73.

108. Yoon H, Triplet EM, Simon WL, Choi CI, Kleppe LS, De Vita E, et al. Blocking Kallikrein 6 promotes developmental myelination. Glia. 2022;70(3):430–50.

109. Yoon H, Radulovic M, Wu J, Blaber SI, Blaber M, Fehlings MG, Scarisbrick IA. Kallikrein 6 signals through PAR1 and PAR2 to promote neuron injury and exacerbate glutamate neurotoxicity. J Neurochem. 2013;127(2):283–98.

110. Patra K, Soosaipillai A, Sando SB, Lauridsen C, Berge G, Møller I, et al. Assessment of kallikrein 6 as a cross-sectional and longitudinal biomarker for Alzheimer’s disease. Alzheimers Res Ther. 2018;10(1):9.

111. Goldhardt O, Warnhoff I, Yakushev I, Begcevic I, Förstl H, Magdolen V, et al. Kallikrein-related peptidases 6 and 10 are elevated in cerebrospinal fluid of patients with Alzheimer’s disease and associated with CSF-TAU and FDG-PET. Transl Neurodegener. 2019;8:25.

112. Diamandis EP, Yousef GM, Petraki C, Soosaipillai AR. Human kallikrein 6 as a biomarker of alzheimer’s disease. Clin Biochem. 2000;33(8):663–7.

113. Pang XW, Chu YH, Zhou LQ, Chen M, You YF, Tang Y, et al. Trem2 deficiency attenuates microglial phagocytosis and autophagic-lysosomal activation in white matter hypoperfusion. J Neurochem. 2023;167(4):489–504.

114. Pan J, Zhang M, Dong L, Ji S, Zhang J, Zhang S, et al. Genome-Scale CRISPR screen identifies LAPTM5 driving lenvatinib resistance in hepatocellular carcinoma. Autophagy. 2023;19(4):1184–98.

115. Jain A, Lamark T, Sjøttem E, Larsen KB, Awuh JA, Øvervatn A, et al. p62/SQSTM1 is a target gene for transcription factor NRF2 and creates a positive feedback loop by inducing antioxidant response element-driven gene transcription. J Biol Chem. 2010;285(29):22576–91.

116. Fei M, Zhang L, Wang H, Zhu Y, Niu W, Tang T, Han Y. Inhibition of Cathepsin S Induces Mitochondrial Apoptosis in Glioblastoma Cell Lines Through Mitochondrial Stress and Autophagosome Accumulation. Front Oncol. 2020;10:516746.

117. Decuypere JP, Parys JB, Bultynck G. ITPRs/inositol 1,4,5-trisphosphate receptors in autophagy: From enemy to ally. Autophagy. 2015;11(10):1944-8.

118. Schultheis N, Jiang M, Selleck SB. Putting the brakes on autophagy: The role of heparan sulfate modified proteins in the balance of anabolic and catabolic pathways and intracellular quality control. Matrix Biol. 2021;100–101:173-81.

119. Wang WT, Han C, Sun YM, Chen ZH, Fang K, Huang W, et al. Activation of the Lysosome-Associated Membrane Protein LAMP5 by DOT1L Serves as a Bodyguard for MLL Fusion Oncoproteins to Evade Degradation in Leukemia. Clin Cancer Res. 2019;25(9):2795–808.

120. Du W, Su QP, Chen Y, Zhu Y, Jiang D, Rong Y, et al. Kinesin 1 Drives Autolysosome Tubulation. Dev Cell. 2016;37(4):326–36.

121. Hovanessian AG. On the discovery of interferon-inducible, double-stranded RNA activated enzymes: the 2’-5’oligoadenylate synthetases and the protein kinase PKR. Cytokine Growth Factor Rev. 2007;18(5-6):351–61.

122. Kim Y, Park J, Kim S, Kim M, Kang MG, Kwak C, et al. PKR Senses Nuclear and Mitochondrial Signals by Interacting with Endogenous Double-Stranded RNAs. Mol Cell. 2018;71(6):1051–63.e6.

123. Costa-Mattioli M, Walter P. The integrated stress response: From mechanism to disease. Science. 2020;368(6489).

