## Supplementary material for "Multiomics approach identifies SERPINB1 as candidate progression biomarker for Spinocerebellar Ataxia type 2": Table S6

**Table S6.** Clinical and molecular characteristics of the extended cohort of patients with SCA2.

| Variables | SCA2 patients  (N=82) | |
| --- | --- | --- |
|  | Range | Mean (SD) |
| AO (yrs) | 15−62 | 39.36 (11.43) |
| DD (yrs) | 2−34 | 13.25 (6.46) |
| SARA score | 3.5−36 | 14.96 (7.43) |
| INAS count | 1−6 | 3.37 (1.45) |
| *ATXN2* expanded alleles (CAG repeat length) | 34−48 | 38.40 (2.83) |
| SERPINB1 (ng/ml) | 0.24−6.74 | 3.15 (1.64) |

AO− age at onset; DD− disease duration; SD− standard deviation
