## Supplementary figures and images for "Multiomics approach identifies SERPINB1 as candidate progression biomarker for Spinocerebellar Ataxia type 2"

### Figure S1

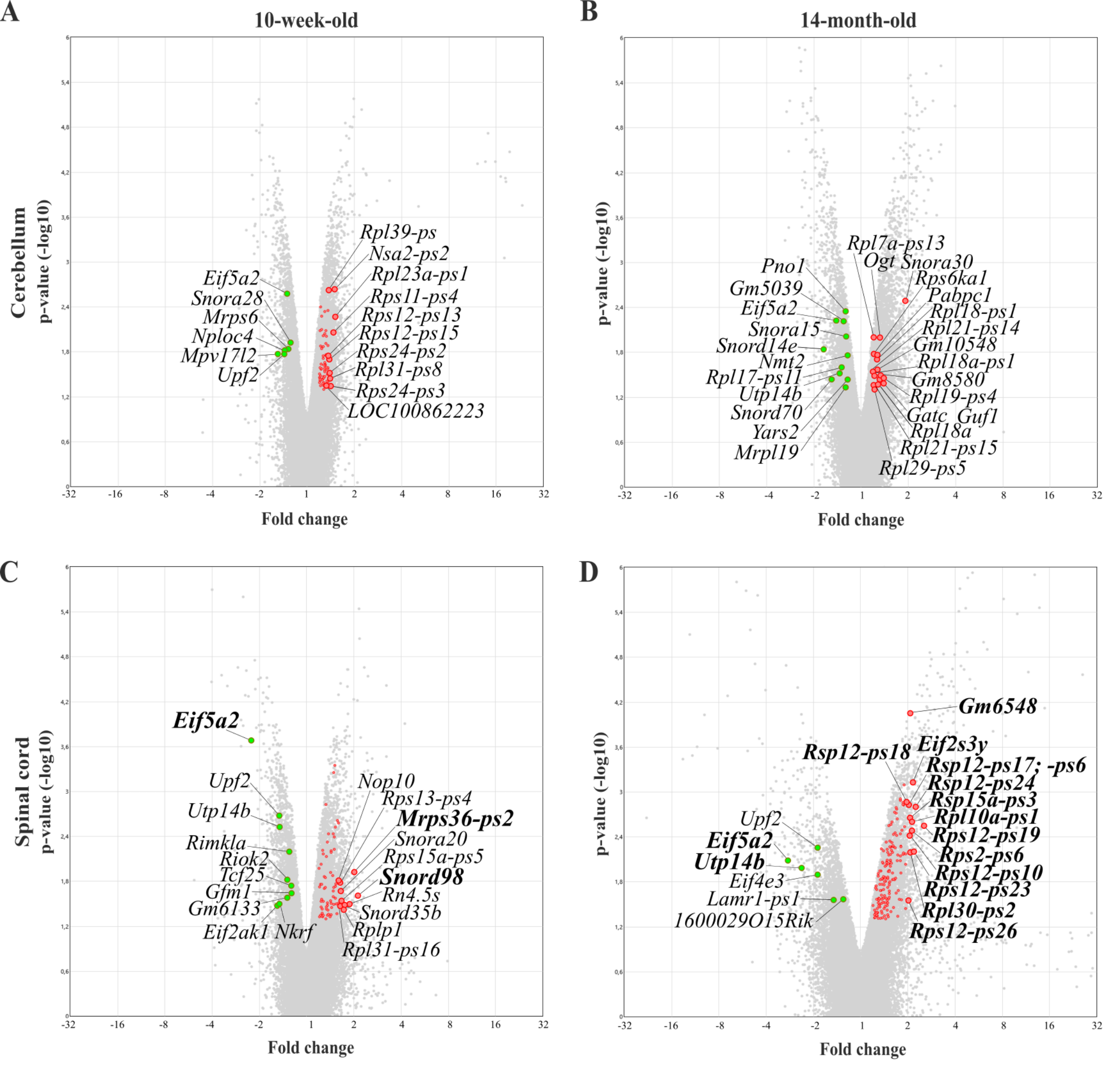

### Figure S2

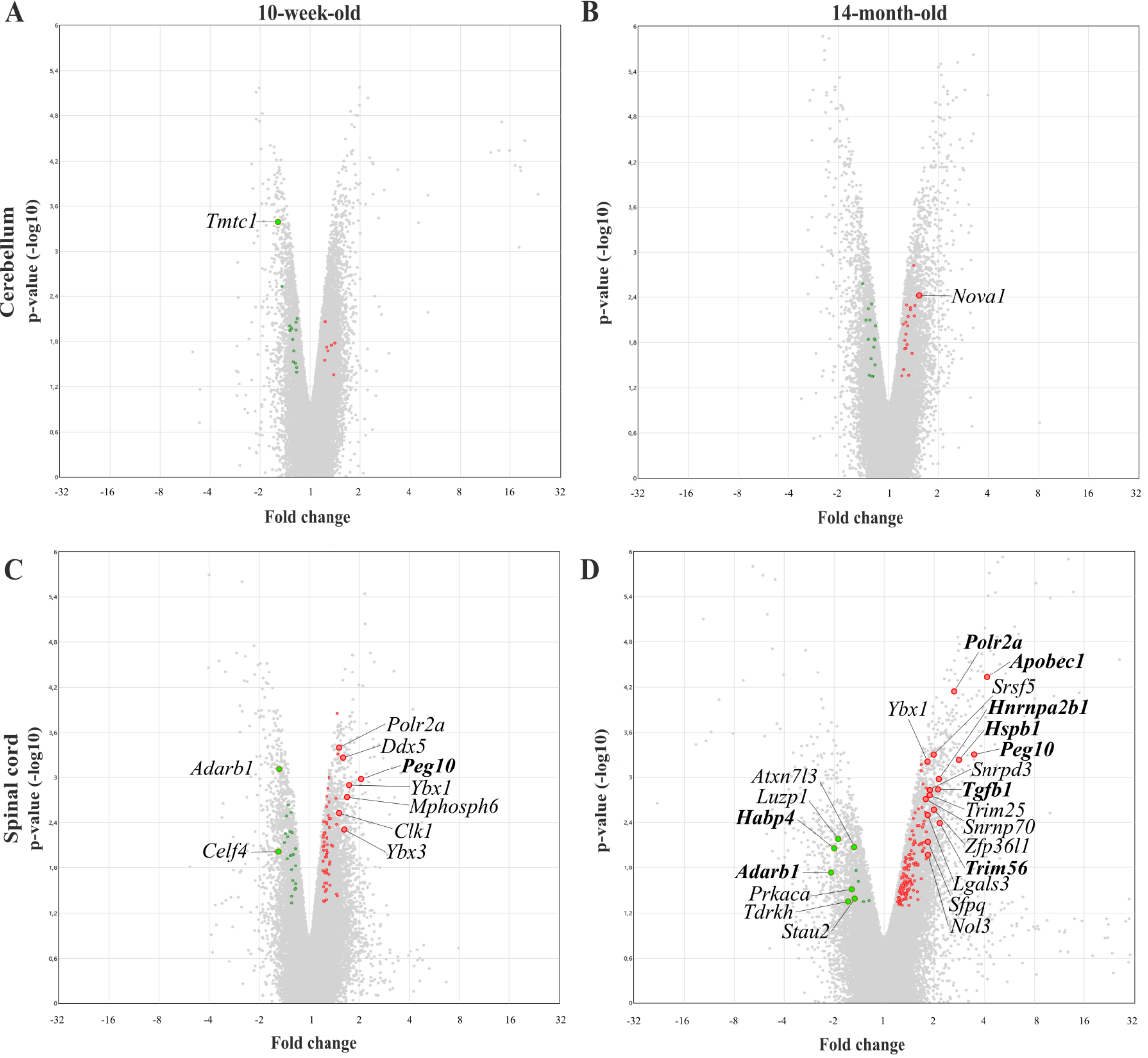

### Figure S3

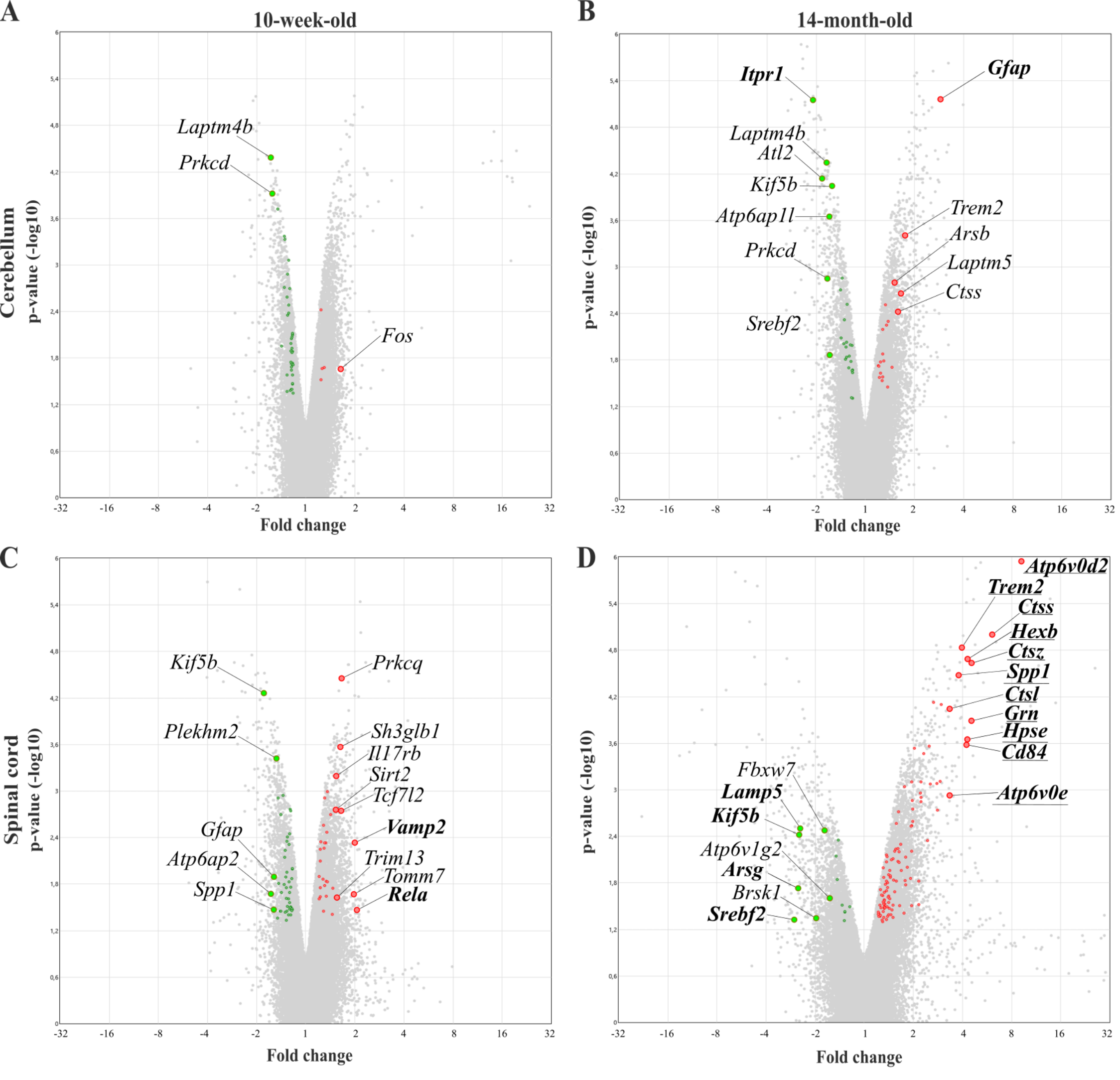

### Figure S4

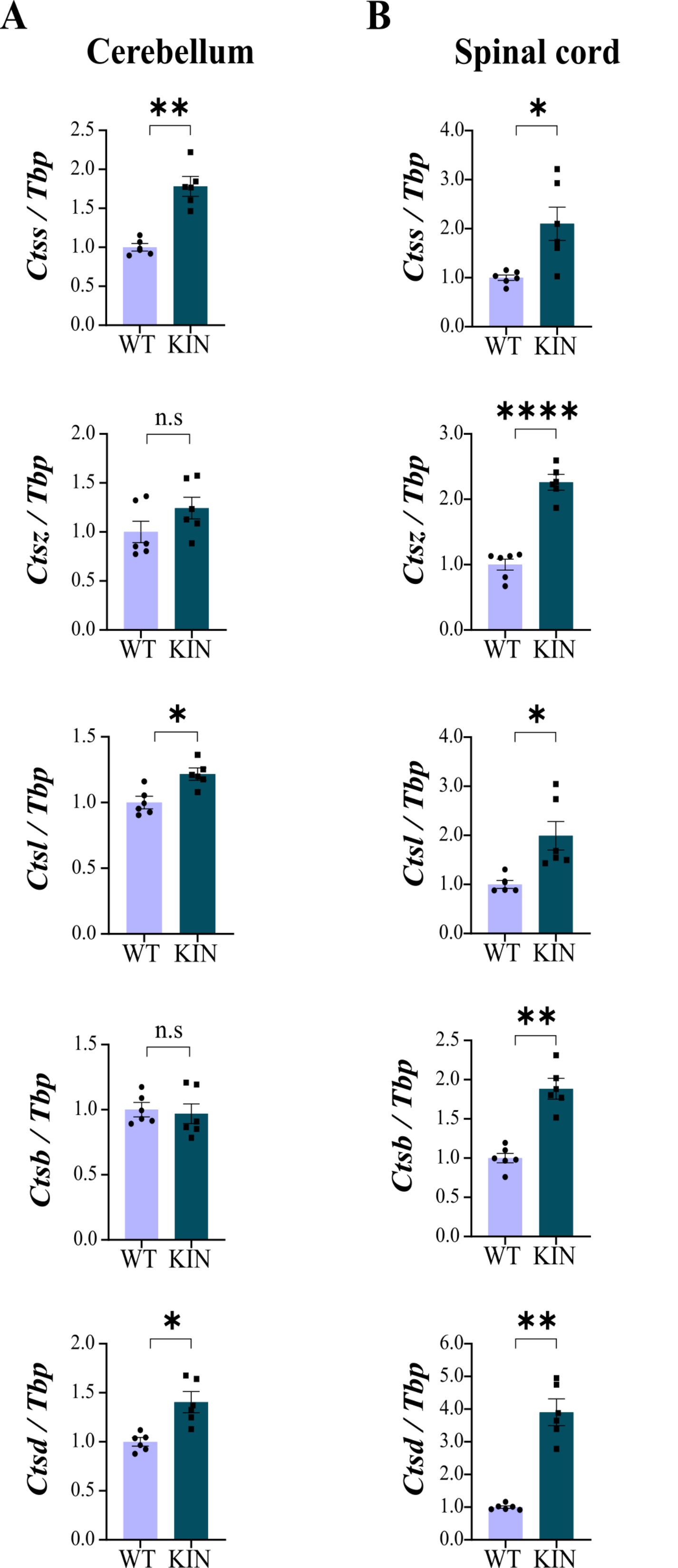

### Figure S5

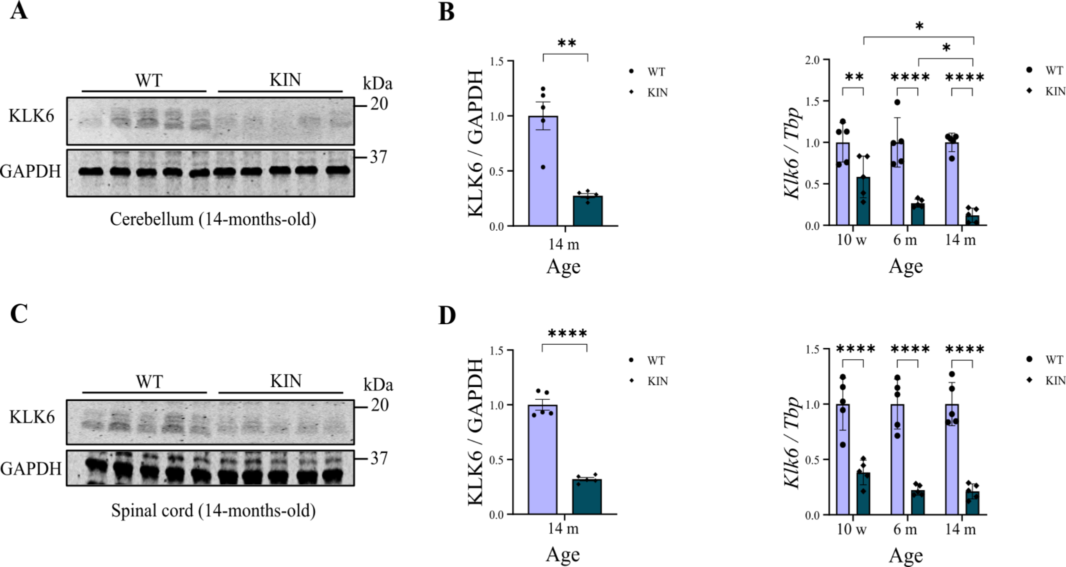

### Figure S6

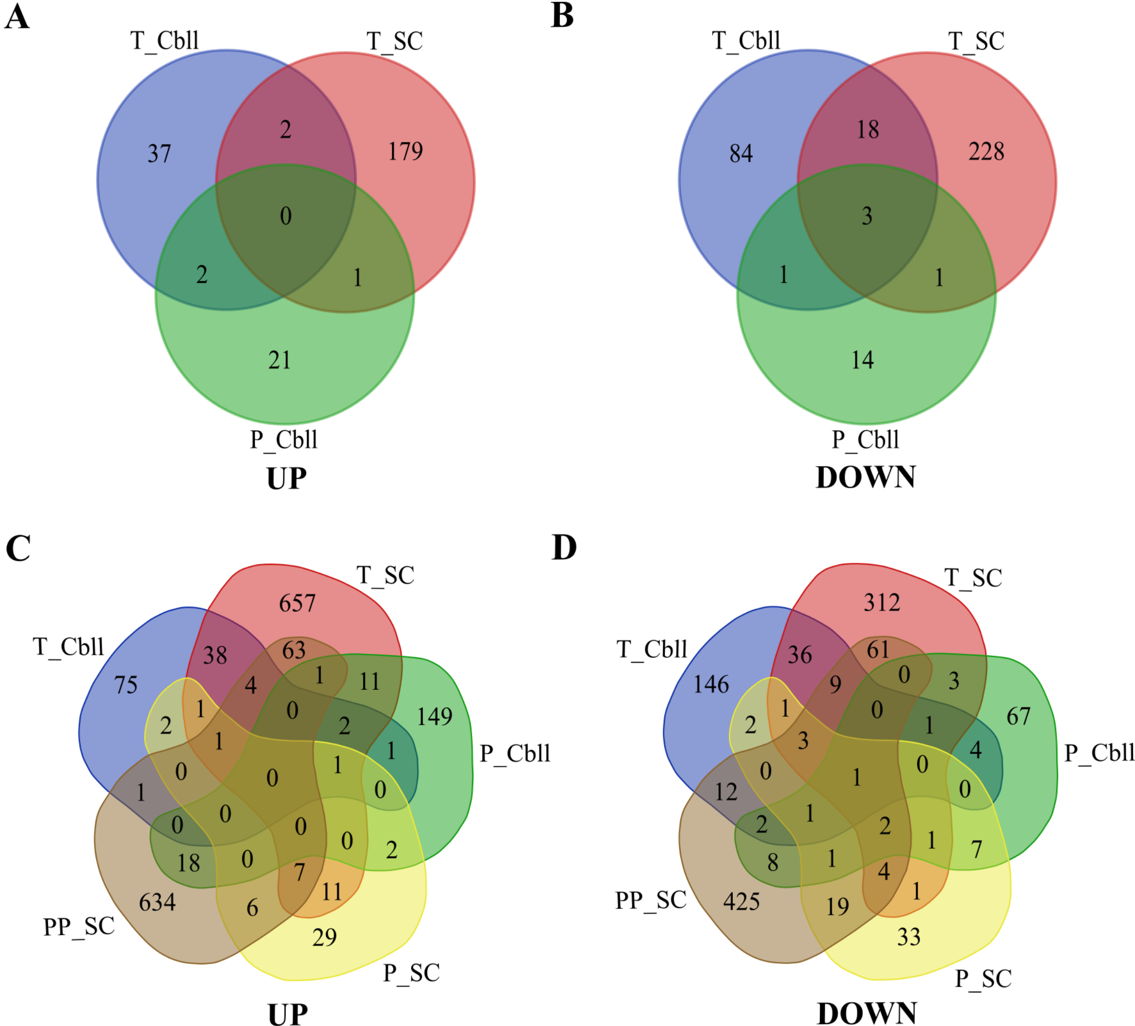

### Figure S7

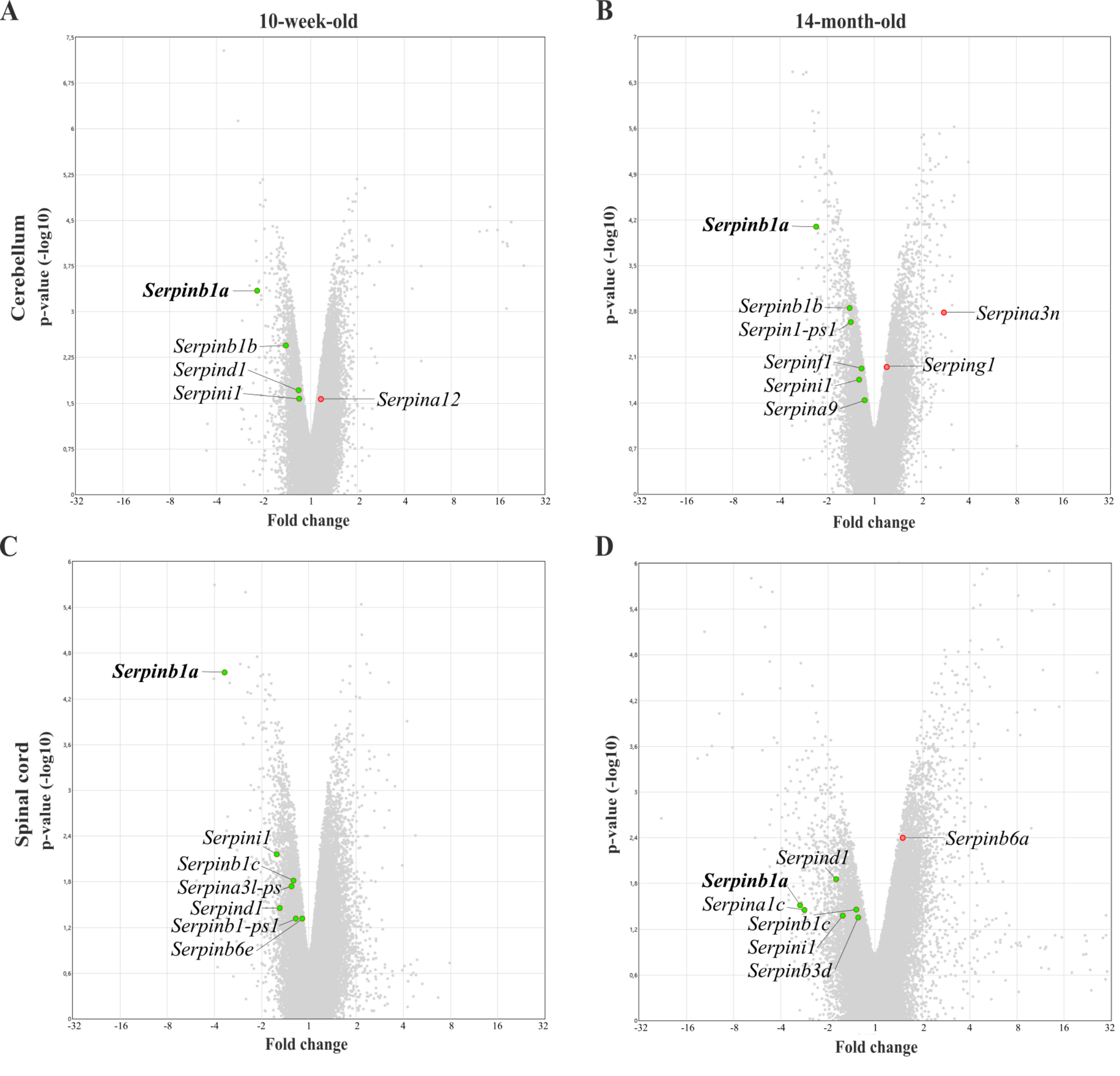

### Figure S8

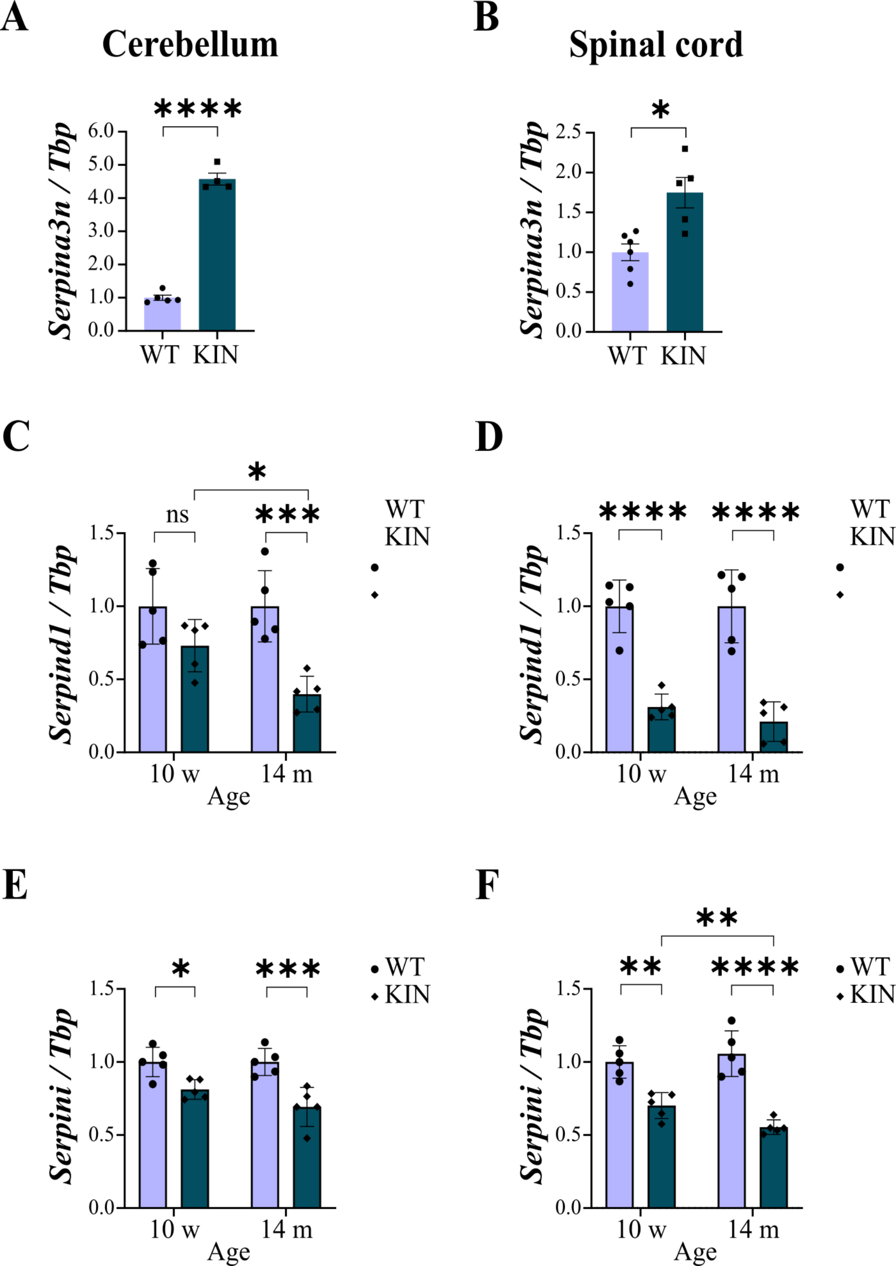
